# Loss of tumor cell MHC Class II drives insensitivity of BRAF-mutant anaplastic thyroid cancers to MAPK inhibitors

**DOI:** 10.1101/2025.01.27.635086

**Authors:** Vera Tiedje, Jillian Greenberg, Tianyue Qin, Soo-Yeon Im, Gnana P. Krishnamoorthy, Laura Boucai, Bin Xu, Jena D. French, Eric J Sherman, Alan L Ho, Elisa de Stanchina, Nicholas D. Socci, Jian Jin, Ronald Ghossein, Jeffrey A. Knauf, Richard P Koche, James A. Fagin

## Abstract

Cancer cells present neoantigens dominantly through MHC class I (MHCI) to drive tumor rejection through cytotoxic CD8+ T-cells. There is growing recognition that a subset of tumors express MHC class II (MHCII), causing recognition of antigens by TCRs of CD4+ T-cells that contribute to the anti-tumor response. We find that mouse Braf^V600E^-driven anaplastic thyroid cancers (ATC) respond markedly to the RAF + MEK inhibitors dabrafenib and trametinib (dab/tram) and that this is associated with upregulation of MhcII in cancer cells and increased CD4+ T-cell infiltration. A subset of recurrent tumors lose MhcII expression due to silencing of *Ciita*, the master transcriptional regulator of MhcII, despite preserved interferon gamma signal transduction, which can be rescued by EZH2 inhibition. Orthotopically-implanted Ciita^-/-^ and H2-Ab1^-/-^ ATC cells into immune competent mice become unresponsive to the MAPK inhibitors. Moreover, depletion of CD4+, but not CD8+ T-cells, also abrogates response to dab/tram. These findings implicate MHCII-driven CD4+ T cell activation as a key determinant of the response of Braf-mutant ATCs to MAPK inhibition.

## Introduction

Antigen presentation, processing and consequent T-cell priming are essential for an effective immune-mediated anti-tumor response. Alterations in the antigen presentation pathway in tumor cells are well described immune escape mechanisms in patients resistant to checkpoint inhibitor therapy (1–3). Intracellular peptides are presented by major histocompatibility complex class I (MHCI) to CD8+ T cells, whereas extracellular peptides are presented by MHC class II (MHCII) molecules to CD4+ T-cells. MHCI is ubiquitously expressed by nucleated cells, whereas MHCII is primarily expressed by professional antigen presenting cells such as dendritic cells (2). Some tumor cell lineages such as breast (4), melanoma (5) and lung (6) express MHCII when exposed to interferon γ (IFNγ). IFNγ leads to JAK1/2 and STAT1 phosphorylation; STAT1 in turn translocates to the nucleus and cooperates with IRF1 to activate the pIV promoter of the gene encoding MHC class II transactivator (CIITA), which functions as a scaffold for the RFX family members RFX5, RFXAP and RFXANK to drive transcription of MHCII-related genes (reviewed in (7)).

Although the presentation of tumor neoantigens through MHCI to CD8+ T-cells is fundamental to the immune response to cancer (8–10), CD4+ T-cells can also eradicate tumors in an antigen-dependent manner in tumor cells that express MHCII (11, 12). Studies in mice implanted with B16 melanoma cells expressing either the Trp1 or pMel model antigens and infused with CD4+ and CD8+ transgenic T-cells that recognize these individually or in combination have helped delineate the contribution of MhcI and MhcII antigen presentation to anti-tumor immune responses (13).

A pooled human kinome shRNA screen of mesothelioma cells identified RET, MAPKK (MEK) and ERK as negative regulators of cell surface abundance of HLA-A02:01, which plays a central role in antigen presentation by MHCI (14), a finding confirmed in other cancer cell lineages. This implicates oncogenic activation of the MAPK pathway in suppressing the antigen presentation machinery driving CD8+ T cell activation. In papillary thyroid cancer (PTC), tumor-cell specific MHCII expression is suppressed in BRAF^V600E^-driven tumors through a TGFβ-dependent autocrine loop (15), and rescued by MAPK pathway inhibitors. BRAF^V600E^ is also the most common MAPK pathway driver in anaplastic thyroid cancer (ATC), commonly associated with mutations of the *TERT* promoter, *TP53* and genes that regulate chromatin remodeling (16, 17). ATC is a devastating disease with a median overall survival that until recently rarely exceeded 6 months. Combined chemoradiation improves survival modestly for stage IVa/b ATC as compared to palliative therapy but provides no benefit to patients with distant metastatic disease (18). A phase II study of *BRAF*^V600E^-mutant ATC treated with the RAF kinase inhibitor dabrafenib and the MEK inhibitor trametinib (dab/tram) showed a remarkable 56% ORR (19, 20). By contrast, the ORR of *BRAF*^V600E^-driven differentiated TC (DTC) to dab/tram was only 30% (21). This was unexpected, as ATCs have a more disrupted genome and aggressive behavior. As compared to DTCs, the tumor microenvironment of ATC is characterized by much heavier infiltration with macrophages and myeloid-derived suppressor cells (MDSCs) and enrichment for NK, CD4^+^ and CD8^+^ T cells (22–24). Overall survival was also improved in ATC patients, however most patients eventually progressed (20, 25). We previously showed in a murine model of ATC driven by thyrocyte-specific dox-inducible BRAF^V600E^ in the context of Tp53 loss that tumors regress profoundly upon BRAF inhibition but often recur. A common mechanism of recurrence is development of *Met* amplification associated with overexpression of its ligand Hgf, resulting in reactivation of MAPK signaling (26). A similar resistance mechanism has been documented in ATC patients with secondary resistance to BRAF/MEK inhibition (27).

Although reactivation of the MAPK pathway is a recognized mechanism of resistance to MAPK pathway inhibitors in several tumor types, the role of tumor cell autonomous expression of components of the antigen presentation machinery in the response to these targeted therapies has not been investigated *in vivo*. This is pertinent to ATC, because their rich inflammatory tumor microenvironment predicts that the response to MAPK inhibitors is in part immune mediated. We found epigenetic silencing of *Ciita* in a subset of recurrent mouse ATC following BRAF inhibition, which led us to investigate the role of MhcII and CD4+ T-cells in the response to dab/tram. We show that *Ciita^-/-^* or *H2-Ab1^-/-^* ATCs are resistant to dab/tram and that CD4+ T-cells are required for the response to these drugs.

## Results

### Human and murine anaplastic thyroid cancer are heavily infiltrated by T-cells and macrophages

We characterized the tumor immune microenvironment (TME) of human PTCs, poorly differentiated thyroid cancers (PDTCs) and ATCs with multiplex immunofluorescence histochemistry on tissue microarrays (Figure 1, A). Among these tumor types, ATCs were the most heavily immune-infiltrated, predominantly with macrophages and T-cells (Figure 1, B). Consistent with this, multispectral flow cytometry of a murine GEMM of ATC (*Tpo-Cre/eYFP/BRaf-CA/Trp53f^l/fl^*; termed *Braf-CA^V600E/^p53*) showed a greater proportion of CD45^+^ cells in relation to live cells than mouse PTC (*Tpo-Cre/Braf-CA^V600E^*; termed *Braf-CA^V600E^*) (Figure 1, C). The greater myeloid population of ATCs as compared to PTCs primarily consisted of macrophages (CD11b^+^ Ly6C^-^ Ly6G^-^ F480^+^) (Figure 1, D), and their lymphoid infiltrate was particularly enriched for FoxP3^-^ CD4^+^ T-cells (Figure 1, E). Further subtyping of the macrophage population showed a more immunosuppressive phenotype in ATCs than PTCs, characterized by arginase-1 (Arg1) and CD206 positive cells with low Mhc class II expression, as well as a higher proportion of PD-L1-expressing macrophages (Figure 1, F).

**Figure 1:**
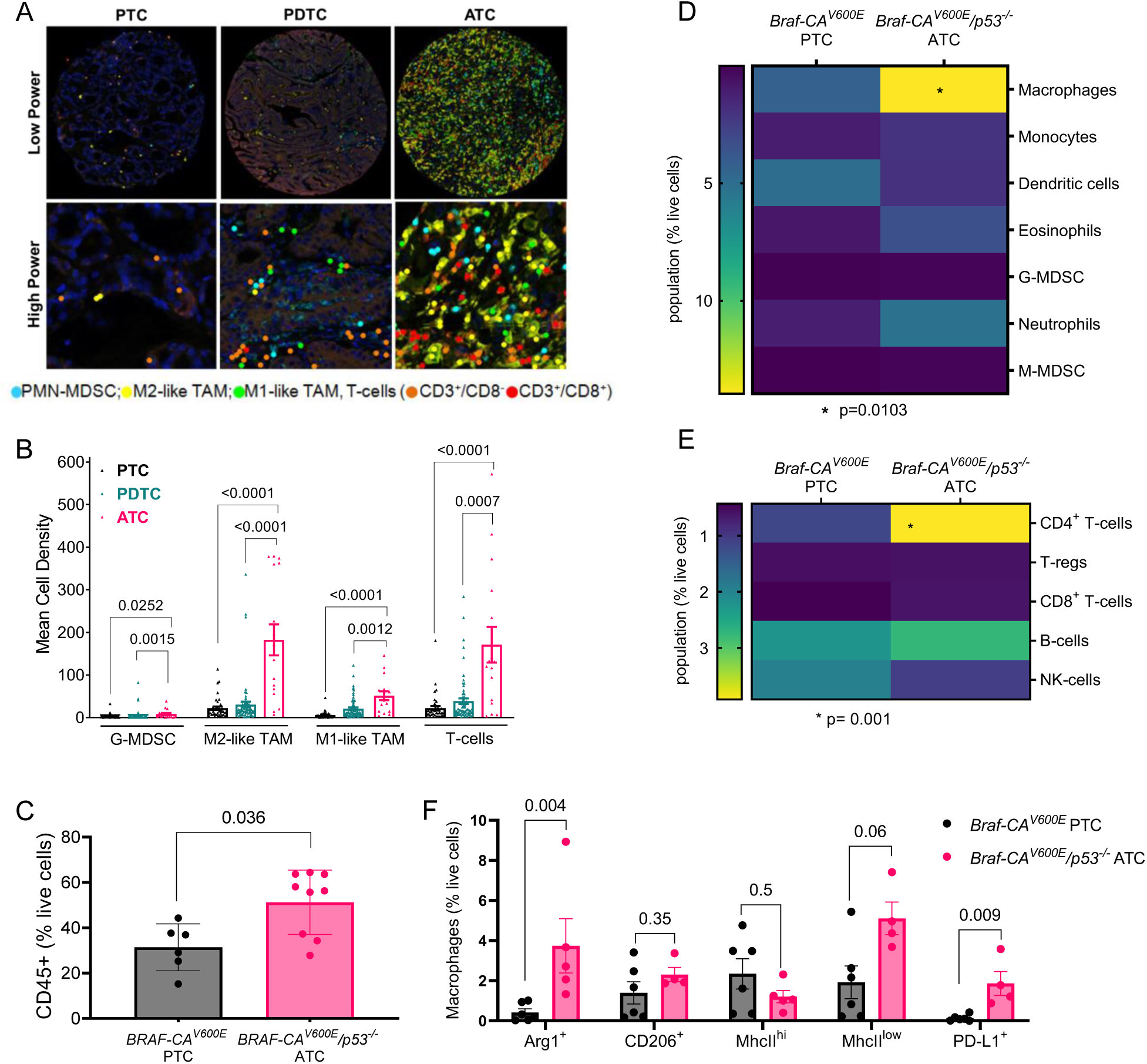
Composition of tumor immune microenvironment in human and murine thyroid cancers. A) Representative images of multiplex immunofluorescence staining for CD3, CD8, CD68, CD163 and CD15 of TMA of 41 PTCs, 72 PDTCs and 16 ATCs. B) Quantification of the TMA for G-MDSC (CD15+), M1-like TAM (CD68+/CD163-), M2-like TAM (CD68+/CD163+) and T-cells (CD3+/CD8- and CD3+/CD8+). C-E) Characterization of immune TME of murine *Braf-CA^V600E^* PTCs (n=6) and *Braf-CA^V600E^/p53^-/-^* ATCs (n=9) by multispectral flow cytometry: C) CD45+ cells. D) Myeloid subpopulations including macrophages (CD11b+ Ly6G-Ly6C-F480+), monocytes (CD11b+ Ly6G-Ly6C+), dendritic cells (CD11c+ F480-MhcII+), eosinophils (CD11b+ Ly6C-Ly6G+ Siglec F+), G-MDSC (CD11b+ Ly6G+ Ly6C-Arg1+), neutrophils (CD11b+ Ly6G+ Ly6C-Arg1-) and M-MDSC (CD11b+ Ly6G-Ly6C+ Arg1+). E) Lymphoid cells including CD4+ T cells (CD3+ CD4+ FoxP3-), T-regs (CD3+ CD4+ FoxP3+), CD8+ T cells (CD3+ CD8+), B-cells (B220+) and NK-cells (B220-NK1.1+). F) Macrophage subtypes. Multiple Mann-Whitney tests (B, C and F); Two-sided ANOVA with Sidak’s multiple comparisons test, with single pooled variance (D, E). Bars represent SEM. TMA: Tissue microarray; PTC: Papillary thyroid cancer; PDTC: Poorly differentiated thyroid cancer; ATC: Anaplastic thyroid cancer; G-MDSC: granulocytic myeloid derived suppressor cells; TAM: tumor associated macrophages; TME: tumor microenvironment; M-MDSC: Monocytic myeloid derived suppressor cells; SEM: Standard error of the mean.

Human and mouse *BRAF^V600E^*-driven ATCs have a higher MAPK transcriptional output than PTCs (26). Moreover, the mutational burden of ATCs as determined by whole exome sequencing is comparable in both species (Supplementary Table 1) (26, 28, 29). Hence, GEMM of ATC closely phenocopy their human counterparts and represent a valid model of the disease.

### MAPK pathway inhibition in mouse *Braf^V600E^/p53^-/-^* ATC tumor cells activates an IFNγ transcriptional output and expression of genes in the antigen presentation pathway

Patients with *BRAF^V600E^* driven ATCs show remarkable structural responses to combined RAF and MEK inhibitors, to the extent that this enables neoadjuvant surgical resection in a subset of patients initially presenting with unresectable disease (25). We used two models to investigate the transcriptional responses of mouse Braf^V600E^-mutant ATCs to inhibition of the driver: 1) Mice with thyroid-specific doxycycline (dox)-inducible expression of BRAF^V600E^ in the context of *Tp53* loss, which included a GFP fluorescent reporter for cell tracking (*Tpo-Cre/LSL-rtTA_GFP/tetO-mycBRAF^V600E^/p53^fl/fl^* (BRAF/p53) (26). 2) Immune competent mice with orthotopic implantation of the *Braf^V600E^/p53^-/-^*cell line TBP3743 (Braf/p53) into the thyroid (30). As previously reported, the BRAF/p53 ATCs showed profound responses to BRAF inhibition (BRAFi) by dox withdrawal (Figure 2, A). Combination therapy with dab/tram also resulted in marked tumor shrinkage of orthotopic ATCs (Figure 2, B).

**Figure 2:**
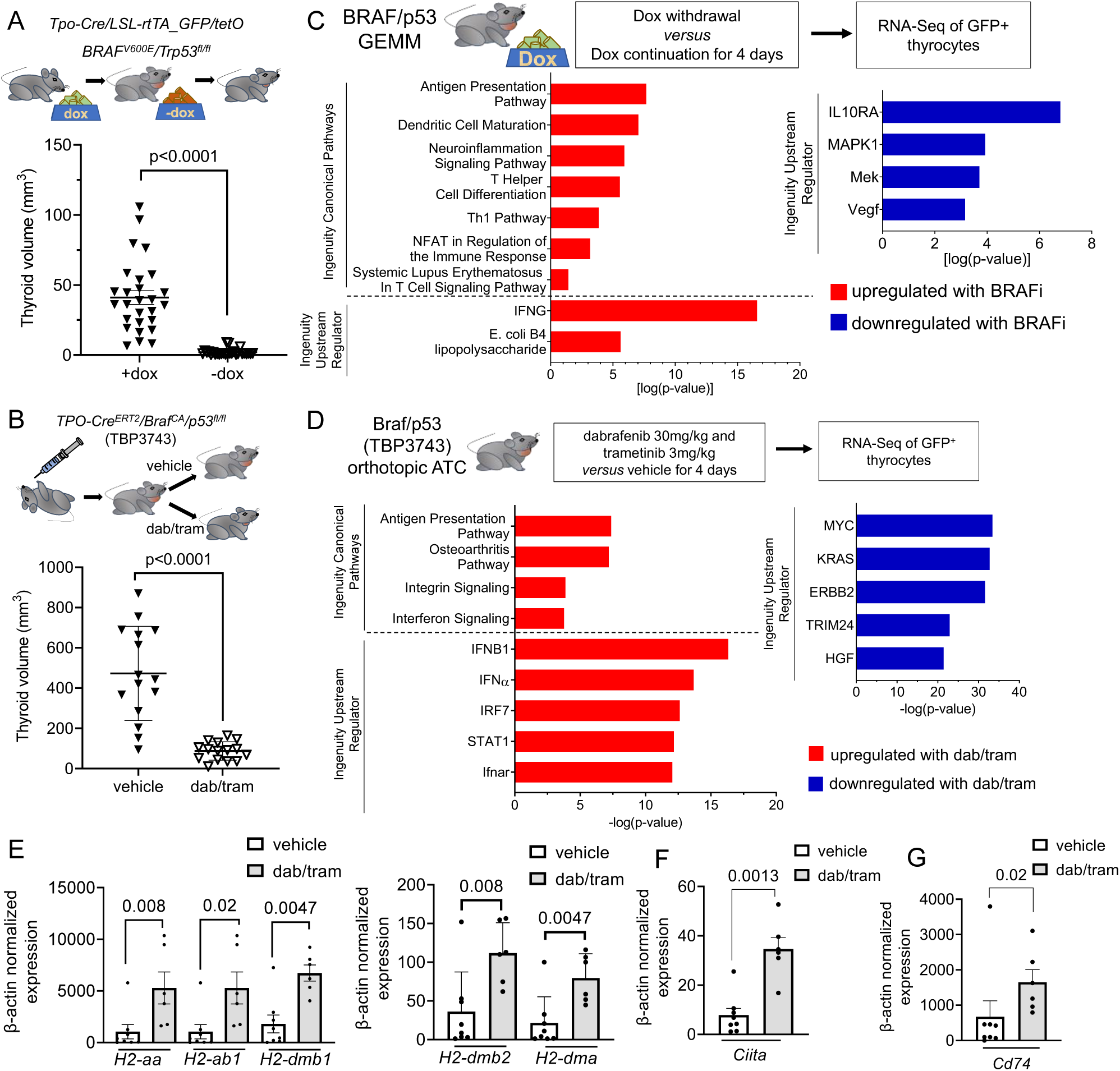
Induction of antigen presentation pathways in BRAF^V600E^ ATC in response to MAPK inhibition. Thyroid volume measurements A) by MRI of dox-inducible BRAF/p53 GEMM on (n=27) and off (n=27) dox for 3-4 weeks and B) by ultrasound in mice orthotopically injected with TBP3743 cells and treated one week after engraftment with dabrafenib 30mg/kg and trametinib 3mg/kg (n=15) or vehicle (n=15) for 2 weeks. Ingenuity pathway analysis of sorted thyrocytes of BRAF/p53 ATCs on and off doxycycline (C) and orthotopic Braf/p53 ATCs treated with vehicle or dab/tram for 4 days (D). E-G) RT-PCR of the indicated MhcII complex mRNAs, Ciita and Cd74 in sorted thyrocytes from Braf/p53 orthotopic ATCs treated with vehicle or dab/tram *in vivo* for 4 days. Multiple Mann-Whitney tests (A, B and E-G). Bars represent SEM. ATC: Anaplastic thyroid cancer; dab/tram: dabrafenib and trametinib; *Ciita*: Class II major histocompatibility complex transactivator; GEMM: Genetic engineered mouse model; SEM: Standard error of the mean.

To understand the transcriptional changes associated with these profound responses to therapy we performed bulk RNA-Seq of FACS-sorted thyrocytes from both models. Ingenuity pathway analysis identified top differentially up- and down-regulated pathways in the context of either BRAF inhibition by dox withdrawal in BRAF/p53 ATCs or dab/tram treatment in the orthotopic Braf/p53 TBP3743 model (Figure 2, C and D). Canonical pathways and upstream regulators enriched by MAPK inhibition in both contexts were related to antigen presentation and interferon pathway activation and, in the BRAF/p53 model, to CD4^+^ T-helper cell maturation.

To further delineate the impact of the MAPK pathway on tumor cell antigen presentation we investigated the expression of genes of the MhcI and II pathways in FACS-sorted thyrocytes from wild type (WT) thyroids and TBP3743 ATCs treated with vehicle or dab/tram for 4 days. MhcI in mice is encoded by the *B2m*, *H2-k1* and *H2-d1* genes, whereas MhcII is encoded by *H2-aa*, *H2-ab1*, *H2-dma*, *H2-dmb1* and *H2-dmb2*. MhcII-related genes were expressed at low levels in orthotopic TBP3743 and BRAF/p53 GEMM ATCs and markedly induced by dox withdrawal or dab/tram treatment, respectively (Figure 2, E, and Supplementary Figure 1, A). Interestingly, MhcII genes are also expressed in wild type thyrocytes, with levels intermediate between those of BRAF/p53 ATCs prior to and after dox withdrawal (Supplementary Figure 1, A). Rfx5, Rfxap and Rfxank are DNA binding proteins that cooperate with Ciita to activate transcription of MhcII-related genes. However, only the expression of *Ciita,* but not that of Rfx genes, was markedly induced by MAPK pathway inhibition (Figure 2, F, and Supplementary Figure 1, B and C). *Cd74*, which is transcriptionally dependent of *Ciita* and assembles with the MhcII alpha and beta chains in the endoplasmic reticulum (ER) to prevent endogenous peptide binding, was suppressed in vehicle treated ATCs and rescued by MAPKi (Figure 2, G, and Supplementary Figure 1, D). By contrast to MhcII genes, expression of the MhcI genes H2-k, H2-d and B2m genes were only modestly suppressed in Braf-ATC vs WT thyroid cells, with expression induced by MAPK inhibition (Supplementary Figure 1, E).

### Recurrent human and murine BRAF-driven ATCs show attenuated MhcII expression in response to MAPK pathway inhibition

To determine whether MHCII expression was associated with response to MAPK inhibitor therapy we performed IHC for HLA-DR in BRAF^V600E^-driven ATC samples from untreated patients, from those showing partial response or stable disease to this therapy and from patients with progressive disease (Figure 3, A; Table 1). HLA-DR expression by ATC cells was dampened in specimens from MAPK inhibitor naïve patients (Figure 3, B). By contrast HLA-DR was expressed by all ATC cells in 4 out of 5 surgical samples from patients with structural response to BRAF and/or MEK inhibitor treatment, whereas significantly fewer ATC cells expressed HLA-DR in specimens from progressive lesions. (Figure 3, C).

**Figure 3:**
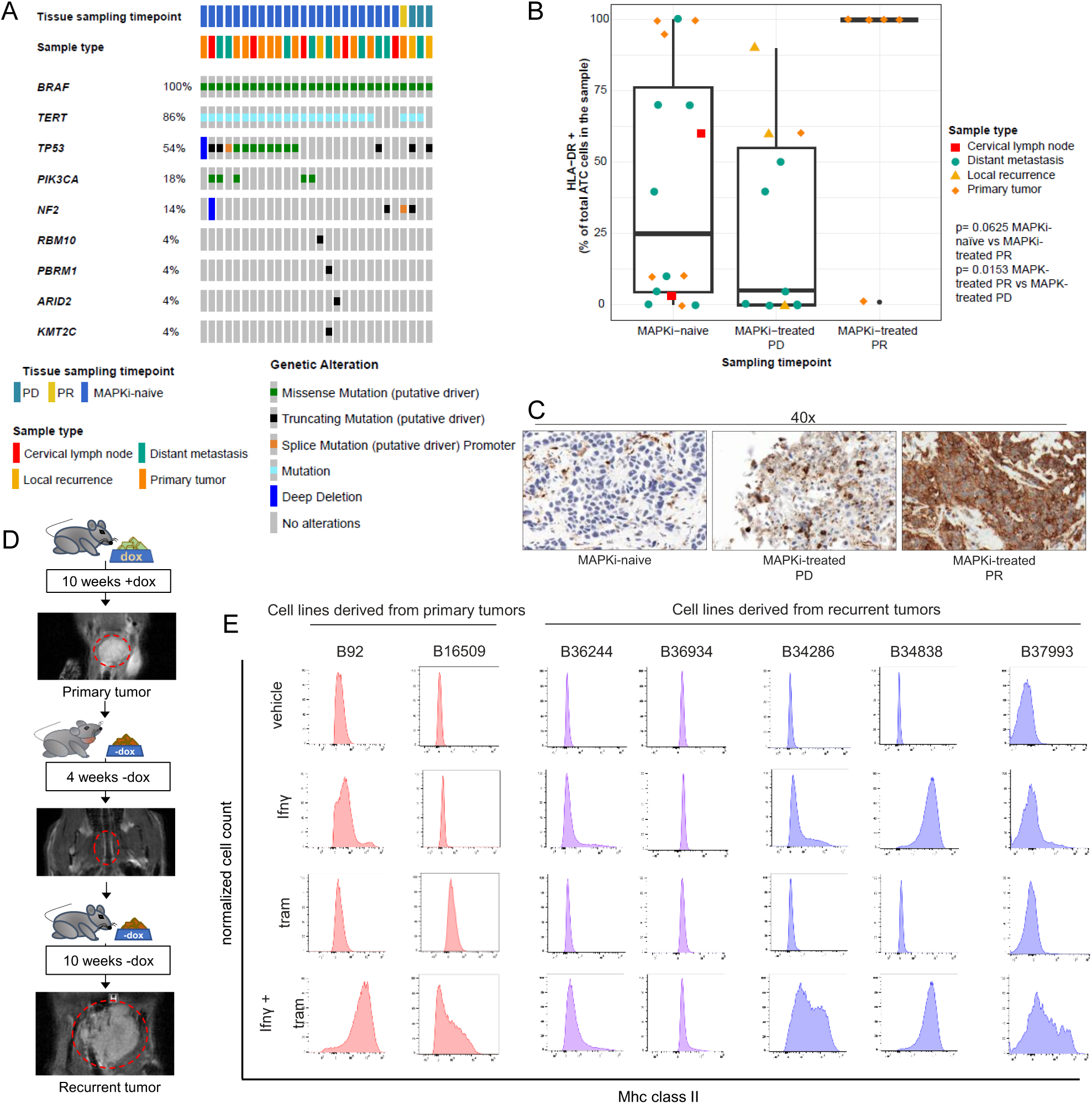
MhcII expression is suppressed in recurrent ATCs. A) Oncoprint of ATC specimens from 28 patients from MSK-clinical cohort sequenced by MSK-IMPACT subjected to HLA-DR IHC. B) Percent of ATC cells expressing HLA-DR in specimens from patients prior to MAPK inhibitor therapy (MAPKi naïve, n=17), at the time of structural response (MAPKi PR, n=5) or disease progression (MAPKi PD, n=10) under therapy with dab/tram (n=22), vemurafenib (n=1), dabrafenib (n=1) or PLX8394 (n=1). C) Representative IHC images of HLA-DR IHC. D) Representative MRIs of a primary ATCs (+dox), the response 4 weeks after dox withdrawal (-dox) and a recurrence at 10 weeks (-dox). E) MhcII flow cytometry of cell lines generated from primary ATCs (B92 and B16509) and recurrent tumors (B36244, B36934, B34286, B34838 and B37993) treated for 96h with vehicle, IFNγ (20ng/ml), trametinib (10nM) or the combination in vitro. Plasma membrane MhcII levels were not induced by IFNγ + tram in the recurrent ATC cell lines B36244 and B36934. Multiple Mann Whitney Tests (B). Mean with SEM. ATC: Anaplastic thyroid cancer; MAPKi: MAPK inhibitor; PR: partial response; PD: progressive disease; dab/tram: dabrafenib and trametinib; IFNγ: Interferon γ. SEM: Standard error of the mean.

**Table 1:** Patient characteristics. n.a.: not applicable

| Patient | Sex | Age at diagnosis | Stage at diagnosis | MAPK-inhibitor treatment regimen | therapy duration [days] |
| --- | --- | --- | --- | --- | --- |
| 1 | F | 56 | IVB | n.a. | n.a. |
| 2 | F | 58 | IVC | dab/tram | 333 |
| 3 | M | 74 | IVC | dab/tram | 268 |
| 4 | M | 63 | IVC | dab/tram | 16 |
| 5 | F | 74 | IVC | n.a. | n.a. |
| 6 | M | 65 | IVB | n.a. | n.a. |
| 7 | F | 66 | IVC | PLX8394 | 644 |
| 8 | M | 58 | IVC | dab/tram | 174 |
| 9 | F | 59 | IVC | dab/tram | 1502 |
| 10 | F | 78 | IVB | dab/tram | 126 |
| 11 | M | 77 | IVC | dab/tram | 81 |
| 12 | F | 58 | IVC | dab/tram | 275 |
| 13 | F | 73 | IVB | dab/tram | 601 |
| 14 | F | 65 | IVC | dab/tram | 310 |
| 15 | F | 68 | IVC | dab/tram | 115 |
| 16 | M | 70 | IVB | dab/tram | 207 |
| 17 | F | 66 | IVC | dab/tram | 1089 |
| 18 | M | 62 | IVC | dab/tram | 229 |
| 19 | M | 78 | IVC | dab/tram | 1166 |
| 20 | F | 58 | IVB | dab/tram | 242 |
| 21 | M | 66 | IVB | vemurafenib | 168 |
| 22 | M | 63 | IVB | dabrafenib | 155 |
| 23 | M | 67 | IVC | dab/tram | 21 |
| 24 | M | 67 | IVC | dab/tram | 191 |
| 25 | F | 78 | IVB | dab/tram | 89 |
| 26 | M | 71 | IVC | dab/tram | 274 |
| 27 | F | 68 | IVC | dab/tram | 109 |
| 28 | M | 71 | IVC | dab/tram | 69 |

Although murine BRAF/p53 ATCs shrank dramatically upon dox-withdrawal, most of them ultimately recurred (Figure 3, D). We previously reported that the recurrent tumors were characterized by BRAF^V600E^-independent mechanisms of MAPK pathway reactivation, such as *Met* gene amplification and *Ras* point mutations, which were detected in a subset of the recurrences (26). We generated cell lines from primary (B92, B16509) and recurrent tumors (B36244, B37993, B34286, B36934, B34838). The B36934 and B34838 lines were derived from *Met*-amplified recurrences.

Since transcriptional induction of antigen presentation pathways, primarily of MhcII, were a key feature of response of mouse ATCs to MAPK inhibition, we next examined whether this response was retained in cell lines derived from recurrent tumors. We determined MhcI and II expression by flow cytometry in two primary and five recurrent cell lines in response to DMSO, IFNγ (20ng/ml), trametinib (10nM) or the combination of IFNγ and trametinib for 96 hours. Whereas IFNγ was sufficient to induce MhcI expression in both primary and recurrent cell lines (Supplementary Figure 2, A), the combination of IFNγ and trametinib was required to maximally induce MhcII in cell lines from the primary ATCs (Figure 3, E and Figure 4, C and D). MhcII expression was not rescued by IFNγ and trametinib in 2 of the 5 recurrent cell lines, B36244 and B36934 (Figure 3, E and Figure 4, A and B). This was not due to impaired inhibition of MAPK by trametinib or of activation of IFNγ downstream signaling (Supplementary Figure 2, B). Notably, expression of *Ciita* and *Cd74* were markedly attenuated in the two recurrent as compared to the two primary cell lines (Figure 4, C and D).

**Figure 4:**
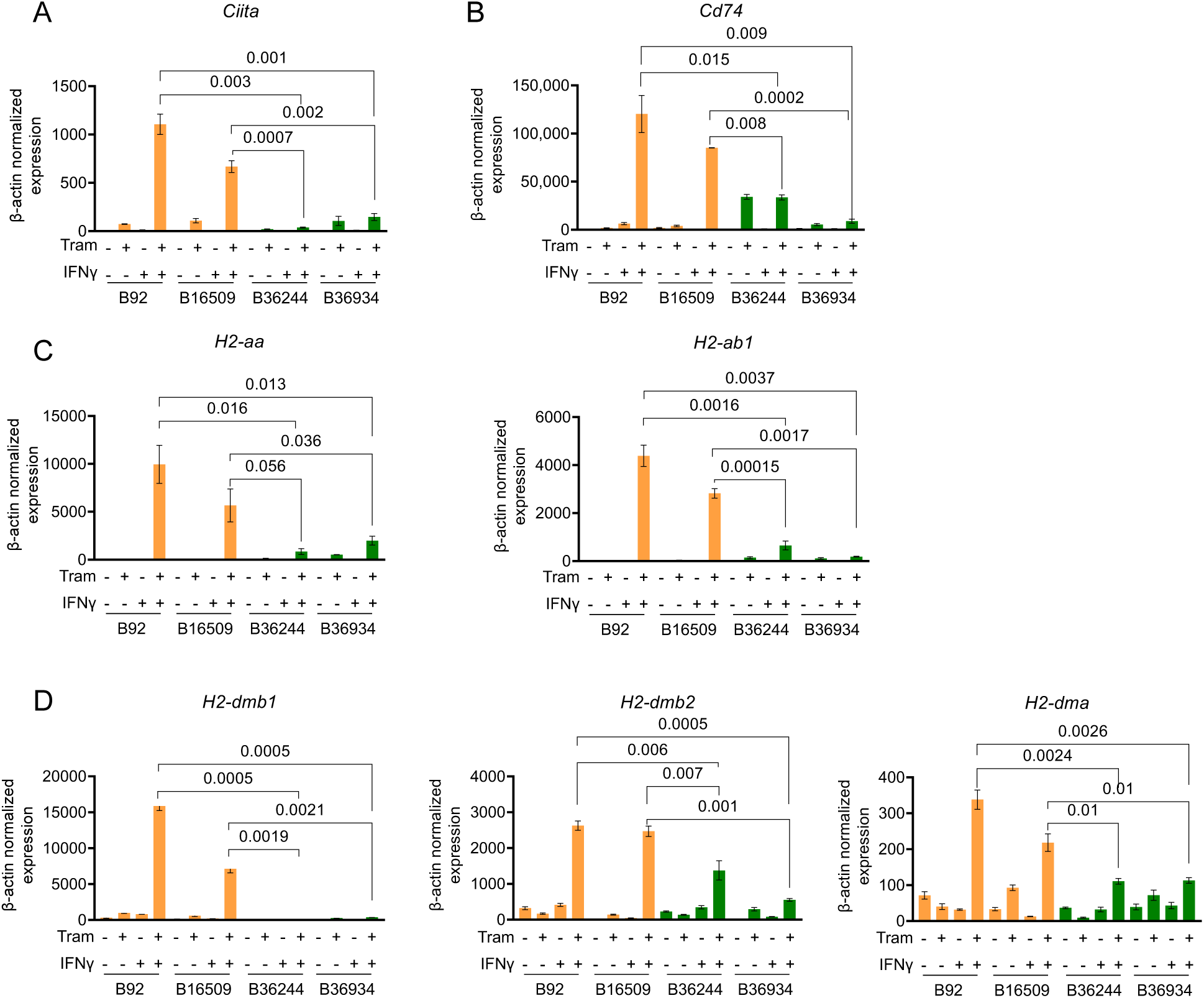
Ciita expression is lost in recurrent ATC cells. A-D) RT-PCR of two primary and two “MhcII-low” recurrent cell lines for the indicated MhcII complex genes (A and B), Ciita (C) and Cd74 (C) in response to IFNγ, trametinib or the combination for 96h. Multiple Mann-Whitney tests. Bars represent SEM. Dox: doxycycline; IFNγ: Interferon γ; Tram: trametinib; Ciita: Class II major histocompatibility complex transactivator; SEM: standard error of the mean.

### Loss of MhcII expression in recurrent ATCs is due to epigenetic silencing of *Ciita*

To probe into the underlying mechanisms of MhcII loss and attenuated *Ciita* expression in recurrent versus primary cell lines we performed bulk RNAseq of a primary (B92) and recurrent (B36934) cell line after a 96h treatment with DMSO, IFNγ (20ng/ml), trametinib (10nM) or the combination of IFNγ and trametinib. It is important to note that these are not isogenic cell lines and that a PCA analysis of their transcriptomes showed that they are distinct from each other at baseline (Supplementary Figure 3, A and B). Because of that we focused our analysis on a previously defined IFNγ transcriptional output (31) and on the genes required for expression of the MhcI and II complexes. Consistent with our flow cytometry data (Figure 3, E), the response to IFNγ was markedly attenuated in both cell lines and rescued by co-treatment with trametinib (Figure 5, A). The MhcII pathway was robustly induced by combination therapy only in the primary cell line (Figure 5, A).

**Figure 5:**
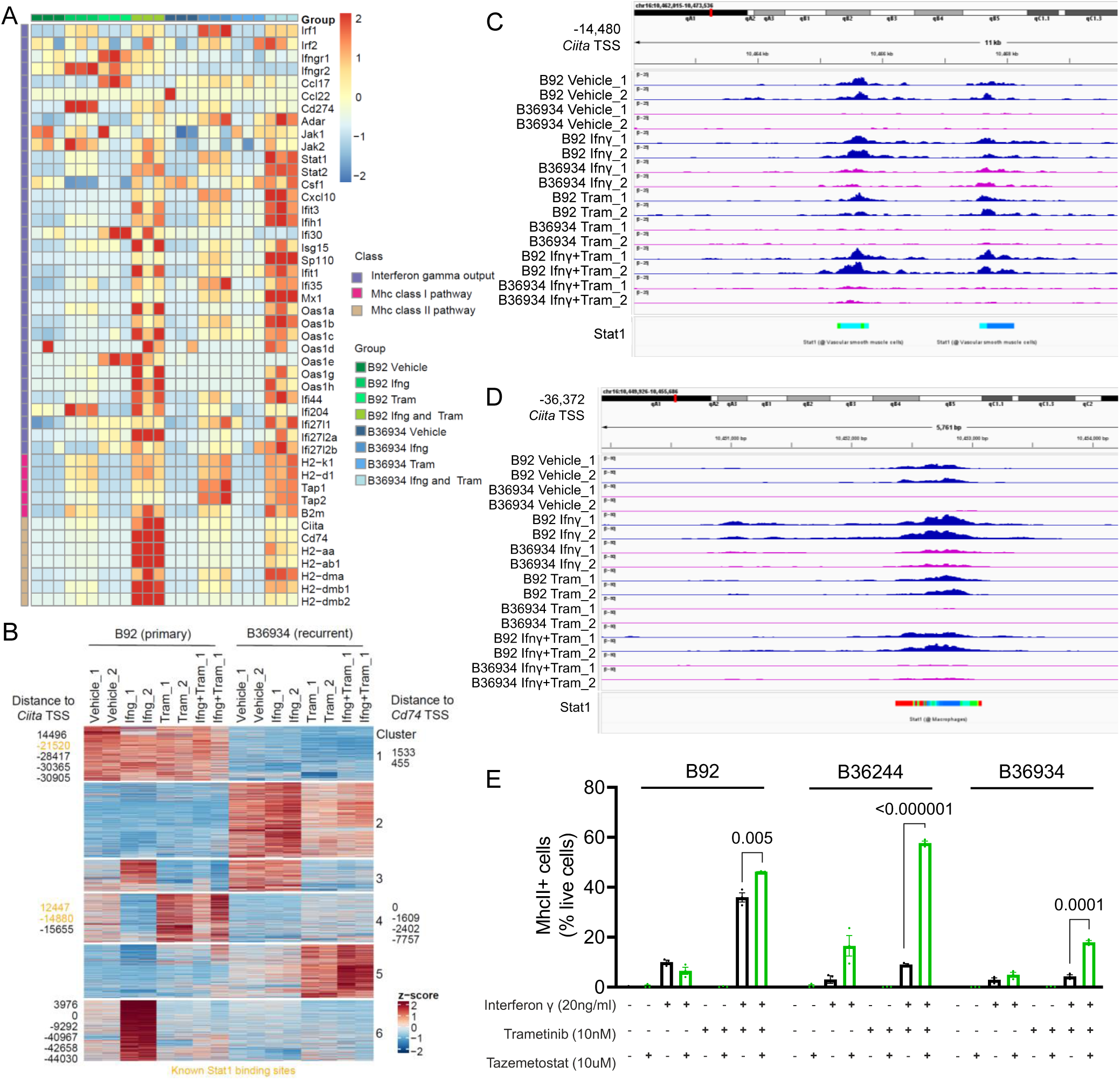
Loss of tumor cell MhcII expression is associated with epigenetic silencing of the Ciita locus. A) RNA Seq of B92 and B36934 cells highlighting MhcI, MhcII and IFNγ-related gene expression pathways in response to a 96h treatment with vehicle, IFNγ, tram or the combination. B) Unsupervised k-means clustering of ATAC-seq peak gains (red) and losses (blue) in B92 and B36934 cells in the indicated treatment conditions. C-D) Integrative Genomics Viewer (IGV) plots showing ATAC-seq peak losses in the recurrent cell line B36934 in comparison to the primary cell line B92 at Stat1 and Irf1 transcription factor binding sites related to the *Ciita* locus. F) Percent of live cells expressing MhcII as determined by FACS in B92 and B36934 cells treated with IFNγ, tram or IFNγ + tram, with or without addition of the EZH2 inhibitor tazemetostat. Ciita: Class II major histocompatibility complex transactivator; IFNγ: Interferon γ. Tram: trametinib.

Since there was no apparent global impairment of IFNγ signaling or transcriptional output between primary and recurrent cell lines (Supplementary Figure 2, B) we tested whether silencing of *Ciita* was associated with decreased chromatin accessibility by ATAC-seq. The ATAC-seq dynamic peaks in the B92 and B36934 lines were resolved into 6 clusters (Figure 5, B). We mapped the peaks that were within 50 kb of the transcription start sites (TSS) of *Ciita* and *Cd74* and found that these were confined to clusters 1, 4 and 6 (Figure 5, B). Chromatin accessibility in these clusters was decreased in the recurrent cell line in all treatment conditions. We integrated our ATAC-seq data with the ChIP Atlas data (32, 33) and found that multiple ATAC peaks around *Ciita* are genuine binding sites for Stat1 and are exclusively accessible in the cell line derived from the primary tumor (Figure 5, C and D). As reflected in the gene expression analysis, global analysis of Stat1 and Irf1 binding sites based on previously documented data in IFNɣ-stimulated macrophages did not show differences between the 6 clusters defined by our ATAC-seq data (32, 33) (Supplementary Figure, 4). We used HOMER motif analysis to identify transcription factor motif enrichments in the 6 ATAC clusters. Motifs related to transcriptional regulators of MhcII genes such as Stat and Rfx-family members are enriched in cluster 1, 4 and 6 (Supplementary Figure 3, C-E).

*Ciita* expression has been reported to be dependent on BRG1 (SMARCA4), one of the two mutually exclusive ATPases that serve as catalytic subunits of SWI/SNF chromatin remodeling complexes. Upon IFNγ stimulation BRG1 displaces EZH2 and SUZ12, key components of the Polycomb Repressor Complex 2 (PRC2), from inter-enhancer regions across the *Ciita* locus (34). As loss-of-function mutations of SWI/SNF genes are common in advanced thyroid cancers (17, 35), we performed whole exome sequencing on B34286 cells, which retained IFNγ and trametinib-induced MhcII expression, and on the two recurrent cells that did not (B36934 and B36244) using tail DNA from the corresponding mouse line as control and found no non-synonymous mutations in any of the Swi/Snf genes (Supplementary Table 2). To test whether PRC2 activity accounts for the repression of MhcII in the two recurrent cell lines we treated them with tazemetostat, a catalytic EZH2 inhibitor. Pretreatment of B92, B36934 and B36244 cells with tazemetostat for 96h, followed by IFNγ + trametinib for a further 48h either completely or partially rescued MhcII in the recurrent lines (Figure 5, E).

### Response to MAPK inhibition in *BRAF^V600E^/p53^-/-^* ATCs requires tumor cell MhcII expression and is CD4+ T-cell dependent

Orthotopically implanted TBP3743 cells become heavily infiltrated by both CD4^+^ and CD8^+^ T-cells in response to dab/tram (Figure 6A and B), but the relative contribution of these two T cell populations to the response to MAPK inhibition is unknown. Because recurrent mouse ATCs can lose MhcII, we asked whether silencing of *Ciita* and MhcII expression was sufficient to induce resistance to dab/tram. For this we generated homozygous CrisprKO clones of *Ciita* and *H2-Ab1*, a major component of the MhcII complex, in TBP3743 cells (Supplementary Figure 4, A, B and F). We confirmed that MhcII expression as measured by FACS was absent in both *Ciita^-/-^* and *H2-Ab1^-/-^* clones (Supplementary Figure 4, C and G). We generated orthotopic models with *Ciita^-/-^* and *H2-Ab1^-/-^* ATCs, treated the mice with dab/tram (schema in Figure 6, C and D) and performed serial ultrasounds to assess therapy response as determined by tumor volume change. Whereas the IC_50_ to trametinib of two independently derived clones of *Ciita^-/-^* and *H2-Ab1^-/-^* was comparable to that of the parental cell line *in vitro* (Supplementary Figure 4, D and H), the *H2-Ab1^-/-^* and *Ciita^-/-^* TBP3743 cells showed attenuated responses to dab/tram *in vivo* (Figure 6, C and D). Moreover, depletion of CD4^+,^ but not CD8^+^ T-cells, impaired the response to dab/tram (Figure 6, E, Supplementary Figure 6, E and F).

**Figure 6:**
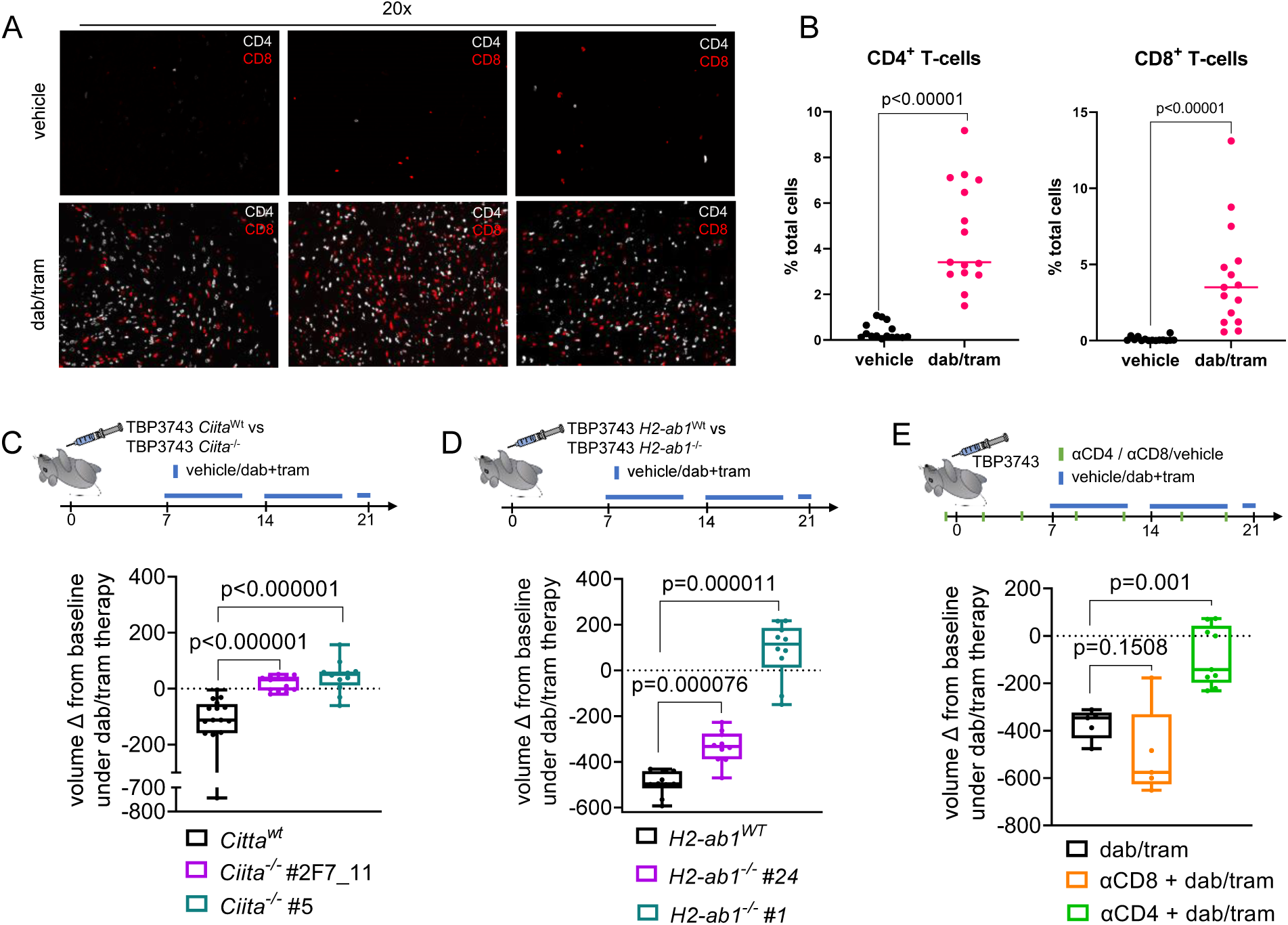
Response of BRAF^V600E^ driven ATCs to dab/tram is lost in Ciita^-/-^ and H2-ab1^-/-^ ATCs and is CD4^+^ T-cell dependent. A) Representative CD4 and CD8 immunofluorescence of Braf/p53 ATCs treated with vehicle or dab/tram for 10 days. B) Quantification of CD4^+^ and CD8^+^ T cells in ATC slides as determined by immunofluorescence. Each dot represents an individual tissue specimen, n=15 for each treatment condition. C) Tumor volume changes of *Ciita^wt^* (n=16) and two clones (n=13 for #2F7_11 and n=12 for #5) of *Ciita^-/-^* TBP3743 cells in response to dab/tram for 2 weeks. D) Response of *H2-ab1^wt^* (n=10) and 2 clones of *H2-ab1^-/-^* (n=10 for #24 and n=11 for #1) to dab/tram treatment for 2 weeks. E) Tumor volume change of orthotopic ATCs in response to treatment with dab/tram (n=9) or in combination with αCD4 (n=6) or αCD8 (n=5) depletion antibody. Multiple Mann Whitney tests (B-E). Bars represent mean with SEM. Whiskers display minimum to maximum. Dab/tram: dabrafenib and trametinib. Ciita: Class II major histocompatibility complex transactivator; ATC: Anaplastic thyroid cancer. SEM: standard error of the mean.

**Figure 7:**
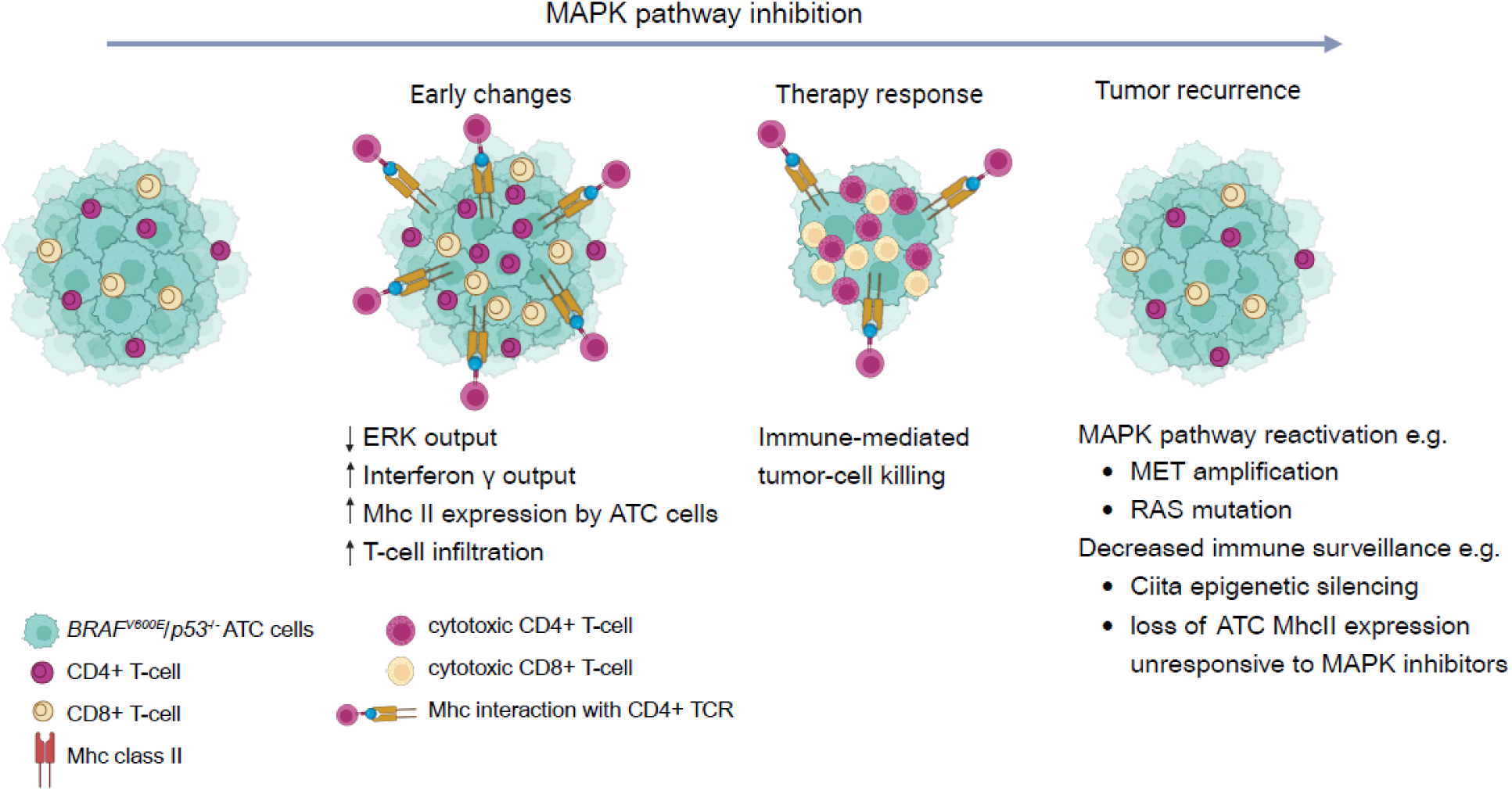
Mechanisms of response and recurrence to MAPK pathway inhibition in BRAF^V600E^/ p53^-/-^ ATC. Upon MAPK inhibition there is an induction of IFNγ transcriptional output in ATC cells leading to MhcII expression and T-cell mediated tumor cell death. Recurrences can occur through MAPK pathway reactivation and/or decreased immune surveillance due to Ciita silencing and loss of MhcII expression.

## Discussion

In this study we show that Braf-mutant ATC tumor cells markedly activate antigen presentation programs in response to therapies targeting the oncoprotein, and that induction of MHCII expression is a particularly determinant factor in response to therapy. The most persuasive evidence supporting this is the discovery that combined treatment of ATC cell lines derived from recurrent tumors with trametinib and IFNγ failed to induce MHCII, and that this is due to epigenetic silencing of *Ciita*. Moreover, knockdown of either *Ciita* or *H2-Ab1*, or depletion of CD4+T cells, is sufficient to abrogate response to MAPK inhibitors *in vivo.* This implicates modulation of the interaction of tumor cells with the immune microenvironment as a key mechanism of response to dabrafenib and trametinib in Braf-mutant ATC. A prior study reported that treatment of *BRAF* mutant PTC cell lines with MAPK inhibitors induced MHC2 expression through a TGFβ autocrine loop (15), but the implications of this finding to resistance mechanisms had not been previously explored.

There are parallels to these findings in BRAF-mutant melanomas. Treatment of human melanoma cell lines with the RAF inhibitor vemurafenib restores expression of melanoma antigens, which are recognized by antigen-specific T cells (36). Moreover, treatment with MAPK inhibitors resulted in marked tumor infiltration with both CD4+ as well as CD8+ T cells after 7 days (37), consistent with our findings in the murine ATC model. Interestingly, a clinical trial of the combination of the MEK inhibitor cobimetinib with vemurafenib showed an increase in proliferating CD4+ T helper cells with no increase in T regulatory cells (38).

Resistance to therapies targeting oncogenic BRAF in murine and human thyroid cancers arise in part due to reactivation of MAPK signaling through mutations of other genes in the pathway, such as *KRAS, HRAS*, *RAC1*, *NF1* or through copy number abnormalities leading to *MET* gene amplification (26, 27, 39). Interestingly, the mouse ATC cell line B36934, derived from a recurrent tumor, harbors a *Met* gene amplification (26) as well as silencing of *Ciita* and refractoriness to induction of Mhc2 by combined treatment with trametinib and IFNγ. Hence, these two distinct mechanisms of resistance can coexist.

There is a relative attenuation of the IFNγ transcriptional output of BRAF-mutant ATC cells, which is rescued by MAPK inhibition in both primary and recurrent cell lines. Although there is some heterogeneity in the expression of individual IFNγ-regulated genes, the most manifest difference between the index primary and recurrent cell lines is the absence of induction of the MhcII mRNA cluster in the latter, rather than a global impairment of the entire IFNγ cistrome. Accessibility of transcription factors to the *Ciita* locus is dependent on the integrity of Swi/Snf chromatin remodeling complexes (34). Indeed, IFNγ-induced *Ciita* transcription is BRG1 (SMARCA4) dependent (34), conferred through eviction of the polycomb repressor complex 2 (PRC2) including its EZH2 and SUZ12 subunits. Dysregulated EZH2 recruitment to Ciita was identified as the underlying mechanism for the absence of MhcII expression in breast cancer cell lines (40). A PRC2 gene expression signature was also shown to be inversely associated with MhcII-related gene expression and T-cell gene signatures in malignant melanoma (41). Moreover, EZH2 inhibition in MhcII negative melanoma cell lines resulted in open chromatin at the IFNγ-inducible promoter pIV of the *Ciita* gene and increased MhcII expression (41). Consistent with this, we also found that EZH2 inhibition led to partial MhcII rescue in ATC cell lines derived from recurrent tumors. Tumor cell MHCII expression has been proposed as a predictor of response to checkpoint inhibitor therapy (5), but prior to our study its loss had not been identified as an acquired resistance mechanism to MAPK inhibitors.

CD8+ T-cells are traditionally considered as the primary cytotoxic effector in cancer. Multiple studies have demonstrated that anti-tumor CD4+ T-cell function is broader than functioning as a helper cell to CD8+ mediated killing (42). They were shown to have direct cytotoxic activity upon recognition of peptides presented by MHCII (11, 43) and to cooperate with inflammatory myeloid cells to induce cell death (13). We demonstrate a central role for CD4+ T-cells in mediating the response to oncogene inhibition in *BRAF^V600E^* driven ATCs, as evidenced by the enrichment of CD4+ T-cells in the TME in response to this treatment and the attenuated response in the context of CD4+ T cell depletion. The loss of tumoral MHCII presentation in recurrent ATCs refractory to BRAF inhibitors squarely implicates T cell cytotoxicity as a central mechanism of response to this therapy, which is nominally directed against a cell autonomous driver of the disease. It also points to a critical role of illegitimate MAPK pathway activation in disabling immune surveillance, a process that may be required for disease pathogenesis.

## Methods

### Patient population

After IRB approval 28 patients with confirmed BRAF^V600E^ anaplastic thyroid cancer (ATC) and available paraffin embedded tissue were selected for this study. Chart review was performed to identify the MAPK inhibitor treatment regimen and therapy response at the time of tissue collection (Table 1). Patients were either treated with dabrafenib and trametinib (n=22), vemurafenib (n=1), dabrafenib (n=1), the pan-RAF inhibitor PLX8394 (n=1) or did not receive MAPK inhibitor therapy (n=3). Specimens collected at different times during the treatment were available from 4 patients. Specimens were either collected prior to MAPK inhibitor treatment (MAPKi naïve), during the treatment with response (MAPKi PR) or with progression (MAPKi PD).

### Multispectral Imaging

MSI was performed by the Vectra 3.0 Automated Quantitative Pathology Imaging System (Perkin Elmer) on tissue microarrays of 41 PTCs, 72 PDTCs and 16 ATCs. Five-micron sections of the tissue microarrays were sequentially stained for CD8, CD3, CD163, CD68 and CD15 on a Bond RX autostainer (Leica). The antibodies used are listed in Supplementary Table 3. Slides were dewaxed and antigen retrieval was performed with epitope retrieval solution 1 or 2 (ER1/2, Leico Biosystems) for 20 minutes at 93°C. Following a 30-minute block (Antibody Diluent reagent; Perkin Elmer), tissues were incubated for 30 minutes with the primary antibody, 10 minutes with horseradish peroxidase (HRP)-conjugated secondary polymer (anti-mouse/anti-rabbit, Perkin Elmer), and 10 minutes with HRP-reactive OPAL fluorescent reagents (Perkin Elmer). Slides were washed between staining steps with Bond Wash (Leica) and stripped between each round (ER1/2). After the final staining, slides were heat-treated (ER1), stained with DAPI (Perkin Elmer), and cover slipped with Prolong Diamond mounting media (Thermo Fisher). Each tissue plug on the microarray was imaged at 20X and analyzed as a single region of interest (ROI). Images were analyzed with inForm software (Perkin Elmer) to unmix fluorochromes, subtract autofluorescence, segment tissue tumor and stromal regions, segment cellular compartments, and phenotype the cells according to morphology and cell marker expression. Cell phenotypes were defined as: CD8+ T cell: CD8+CD3+; “M1-like” macrophage: CD68+CD163-; “M2-like” macrophage: CD68+CD163+; and PMN-MDSC: CD15+

### HLA-DR immunohistochemistry

Slides were incubated with HLA-DR antibody (Supplementary Table 3) at a dilution of 1:500 for 30 minutes. The percentage of tumor cells showing membranous immunopositivity of HLA-DR was recorded.

### ATC mouse models

We generated a murine ATC model by crossing *Tpo-Cre* (PMID: 15282748), *LSL-eYFP* (Jackson Laboratory; stock number 007903), *BRaf-CA* (PMID: 17299132) (Jackson Laboratory; stock number: 017837) and *Trp53^fl,fl^*(PMID: 10783170) (Jackson Laboratory; stock number: 008462) mice to create quadruple *Tpo-Cre/eYFP/BRaf-CA/Trp53^fl/fl^* transgenics, herein referred as Braf-CA^V600E^/p53 ATC, all alleles were backcrossed into the C57BL/6J background (Jackson Laboratory; stock number: 000664). These multi-transgenic mice result in Tpo-Cre driven thyroid-specific expression of BrafV600E, loss of p53 and expression of YFP. Additionally, we generated *Tpo-Cre/eYFP/BRaf-CA* that served as a PTC control (Braf-CA^V600E^ PTC). The *Tpo-Cre/Lsl-rtTA_GFP/tetO-mycBRAFV600E/Trp53^fl/f^* GEMM referred as BRAF/p53 in this manuscript was previously described (26). For the induction of thyroid-specific expression of BRAFV600E mice were fed dox-impregnated chow (2,500 ppm, Envigo). Orthotopic ATCs (Braf/p53) were generated by ultrasound guided injection of 5ul PBS containing 50,000 TBP3743 cells (30) into the right thyroid lobe of F1 B6129SF1/J mice, purchased from Jackson Laboratory. Briefly, mice were anesthetized by inhalation of ∼2% isoflurane with ∼2% O2 and neck hair was removed using defoliating agent. Orthotopic tumor growth was monitored by weekly ultrasound.

### Mouse imaging studies

For ultrasound imaging mice were anesthetized by inhalation of 1.5– 2.5% isoflurane with 2% O_2_. An aqueous ultrasonic gel was applied on the denuded neck and thyroid tumors were imaged with the VisualSonics Vevo™ 770 In Vivo High-Resolution Micro-Imaging System (VisualSonics Inc, Toronto, Ontario, Canada). Using the Vevo™ 770 scan module, the entire thyroid bed was imaged with captures every 250 microns. Using the instrument’s software, the volume was calculated by manually tracing the margin of the tumor every 250 microns. MRI of thyroid tumors from the *Tpo-Cre/LSL-rtTA_GFP/tetO-mycBRAF^V600E^ Trp53^fl/fl^* genetic engineered mice was done as described (26).

### *In vivo* drug studies

Treatment with 30mg/kg dabrafenib (Selleck Chemical) and 3mg/kg trametinib (Selleck Chemical) Monday to Friday via oral gavage was initiated at least 7 days after orthotopic injection of the TBP3743 cell line. For CD4+ and CD8+ T-cell depletion studies, treatment was initiated one day prior to tumor implantation with 200μg of either αCD4 (clone GK1.5, BioXCell) or αCD8 (clone 2.43, BioXCell) and consequently 3x/week for the duration of the study. Tumor growth was assessed by serial ultrasounds.

### Mouse H&E and Immunofluorescence

Mice were euthanized with CO_2_ according with institutional guidelines. Thyroid tumors were harvested and fixed in 4% paraformaldehyde, embedded in paraffin, sectioned and stained with hematoxylin and eosin (H&E) by the MSK Molecular Cytology Core Facility. Automated multiplex IF was conducted with the Leica Bond BX staining system. Paraffin-embedded tissues were sectioned at 5 μm and baked at 58°C for 1 hr. Slides were loaded in Leica Bond and IF staining was performed as follows: Samples were dewaxed and pretreated with EDTA-based epitope retrieval ER2 solution (Leica, AR9640) for 20 min at 100°C. The multiplex antibody staining, and detection were conducted sequentially. The antibodies are listed in Supplementary Table 3. After each round of IF staining, epitope retrieval was performed for denaturation of primary and secondary antibodies before another primary antibody was applied. Finally, slides were washed in PBS and incubated in 5 μg/ml 4’,6-diamidino-2-phenylindole (DAPI) (Sigma Aldrich) in PBS for 5 min, rinsed in PBS, and mounted in Mowiol 4–88 (Calbiochem). Slides were kept at −20°C.

### Tissue processing for flow cytometry

Mice were euthanized, and thyroid tumors harvested on ice, chopped with razor blades and incubated in 5ml FACS Buffer (1xHPSS, 5% FBS) with 1.5mg/ml Collagenase A (Roche) and 0.6mg/ml bovine DNAse (Sigma Aldrich) at 37^°^C for 45 minutes mixing at 200rcf. Cell suspension is then filtered through 70µM cell strainers (Falcon) and red blood cell (RBC) lysis (10x RBC lysis buffer, Biolegend).

### Flow cytometry

Single cell suspensions 1×10^6^ cell were stained with fixable live and dead stain (Fixable Live and Dead Blue, Thermo Fisher) in PBS, incubated with Fc Block (CD16/CD32 antibody, NovusBio) followed by extracellular antibody cocktail incubation in Brilliant Stain Buffer (BD Biosciences). Cells were then prepared for intracellular stain by incubation with Foxp3 / Transcription Factor Staining Buffer (Tonbo Biosciences) for 1h. Afterwards cells were washed and incubated with the intracellular antibody cocktail for 15 min. Samples were then washed three times in FACS Buffer and subjected to analysis at the 5-L Cytek Aurora instrument. Extracellular and intracellular antibodies are listed in Supplemental Table 3. Analysis was performed using FlowJo Version 10.10.7.

### Cell lines

The TBP3743 cell line was a kind gift from Dr. Sareh Parangi (30). TBP3743 cells were cultured in DME HG 5% fetal bovine serum (FBS, Omega Scientific) and 1% pen/strep/glutamine (PSG, Gemini Bio Products). B92, B16509, B36244, B36934, B34286 and B34838 were generated as previously described (26) and maintained in F12 Coon’s with 5% FBS and 1% PSG.

### Generation of *Ciita* and *H2-ab1* CrisprKO clones

For each Ciita- and H2-ab1 CRISPR-Cas9 knockout, clones were created using a guide RNAs targeting 2 distinct sites in exon 10 for Ciita and exon 2 for *H2-ab1*. Gene-targeting dual-guide RNA with Cas9 and mCherry coexpression vectors for each respective KO were custom designed and synthesized by VectorBuilder Inc. (Supplementary Table 4). The Ciita and H2-ab1 CRISPR-Cas9 plasmids were transfected in TBP3743 cells with Lipofectamine 3000 (Invitrogen). Thirty-six hours after transfection, cells were FACS sorted based on positive mCherry fluorescence, and single-cell clones were isolated, and the gRNA-targeted region was screened by PCR (Supplemental Table 5) to confirm CRISPR knockouts.

### Trametinib dose-response curves

Parental, Ciita-/- or H2-Ab1-/- TBP3743 cells were plated in Ultra-Low Adherence 96-well plates and cultured with increasing concentrations of trametinib for 5 days in 5% FBS, 1% Penicillin/Streptomycin/Glutamine and 0.5% methylcellulose. Tumor cell spheroids were then incubated with CellTiter-Glo® 3D Cell Viability Assay reagents and quantified in a Promega GloMax® 96 Microplate Luminometer. Absolute viability values were converted to percentage viability as compared to DMSO-treated controls. IC_50_ curves were generated with GraphPad Prism V10.0 using non-linear fit of log (inhibitor) vs response (three parameters).

### Western blotting

Cells were lysed in 1 x RIPA buffer (Millipore) supplemented with protease (Roche) and phosphatase inhibitor cocktails I and II (Sigma). Protein concentrations were estimated by BCA kit (Thermo Scientific) on a microplate reader (SpectraMax M5); comparable amounts of proteins were subjected to SDS-PAGE using NuPAGE 4%–12% Bis–Tris gradient gels (Invitrogen) and transferred to nitrocellulose membranes. After overnight application of the primary antibody membranes were incubated with secondary antibodies coupled to horseradish peroxidase (HRP) or IRDye fluorophores for 1 h at room temperature. ERK and STAT blots were imaged using iBright CL1000 (Thermo Fisher Scientific). For EZH2, H3 and H3K37Me3 imaging was done using the LI-COR Odyssey. Western blot antibodies are listed in Supplementary Table 3.

### Real time PCR

One microgram of RNA was subjected to DNase I (Invitrogen) treatment and reverse transcribed using SuperScript III Reverse Transcriptase (Invitrogen) following the manufacturer’s protocol. cDNA was diluted at 1:10 and 2ul used as a template for RT-PCR reactions performed using the Power SYBR Green PCR Master Mix (Applied Biosystems) on QuantStudio 8 pro (Applied Biosystems). For gene expression quantifications the Ct values of the target genes were normalized to βactin. The PCR primers are listed in Supplementary Table 5.

### Bulk transcriptomic sequencing

For sorted tumor cells from the GEMM, 1-2 ng total RNA quantified with RiboGreen with RNA integrity numbers ranging from 7.2 to 10 underwent amplification using the SMART-Seq v4 Ultra Low Input RNA Kit (Clonetech catalog # 63488), with 12 cycles of amplification. Subsequently, 9-10 ng of amplified cDNA was used to prepare libraries with the KAPA Hyper Prep Kit (Kapa Biosystems KK8504) using 8 cycles of PCR. Samples were barcoded and run on a HiSeq 4000 in a PE50 run, using the HiSeq 3000/4000 SBS Kit (Illumina). An average of 53 million paired reads were generated per sample and the percent of mRNA bases per sample ranged from 65% to 78% and ribosomal reads averaged 1%. For the sorted tumor cells from orthotopic ATCs 1 µg of total RNA with DV200 percentages varying from 49-90% underwent ribosomal depletion and library preparation using the TruSeq Stranded Total RNA LT Kit (Illumina catalog # RS-122-1202) according to instructions provided by the manufacturer with 8 cycles of PCR. Samples were barcoded and run on a NovaSeq 6000 in a PE100 run, using the NovaSeq 6000 S4 Reagent Kit (200 Cycles) (Illumina). On average, 34 million paired reads were generated per sample and 60% of the data mapped to the transcriptome.

### ATAC sequencing

Profiling of chromatin was performed by ATAC-Seq as described (44). Briefly, ∼50,000 cells were washed in cold PBS and lysed. The transposition reaction containing TDE1 Tagment DNA Enzyme (Illumina catalog # 20034198) was incubated at 37°C for 30 minutes. The DNA was cleaned with the MinElute PCR Purification Kit (QIAGEN catalog # 28004) and material amplified for 5 cycles using NEBNext High-Fidelity 2X PCR Master Mix (New England Biolabs catalog # M0541L). After evaluation by real-time PCR, 9-11 additional PCR cycles were done. The final product was cleaned by aMPure XP beads (Beckman Coulter catalog # A63882) at a 1X ratio, and size selection was performed at a 0.5X ratio. Libraries were sequenced on a NovaSeq 6000 in a PE100 run, using the NovaSeq 6000 S4 Reagent Kit (200 Cycles) (Illumina). An average of 36 million paired reads were generated per sample.

### Whole exome capture and sequencing

After PicoGreen quantification and quality control by Agilent BioAnalyzer, 100 ng of DNA were used to prepare libraries using the KAPA Hyper Prep Kit (Kapa Biosystems KK8504) with 8 cycles of PCR. After sample barcoding, 340-500 ng of library were captured by hybridization using the SinglePlex Mouse Exome (Twist catalog # 102036) according to the manufacturer’s protocol. PCR amplification of the post-capture libraries was carried out for 12 cycles. Samples were run on a NovaSeq 6000 in a PE100 run, using the NovaSeq 6000 S4 Reagent Kit (200 Cycles) (Illumina). Depth of sequencing averaged 78X.

### Bioinformatic analysis for WES

Illumina (HiSeq) Exome Variant Detection Pipeline: The data processing pipeline for detecting variants in Illumina HiSeq data is as follows. First the FASTQ files are processed to remove any adapter sequences at the end of the reads using cutadapt (v1.6). The files are then mapped using the BWA mapper (bwa mem v0.7.12). After mapping the SAM files are sorted and read group tags are added using the PICARD tools. After sorting in coordinate order, the BAM’s are processed with PICARD MarkDuplicates. The marked BAM files are then processed using the GATK toolkit (v 3.2) according to the best practices for tumor normal pairs. They are first realigned using ABRA (v 0.92) and then the base quality values are recalibrated with the BaseQRecalibrator. Somatic variants are then called in the processed BAMs using muTect (v1.1.7) for SNV and the Haplotype caller from GATK with a custom post-processing script to call somatic indels. The full pipeline is available at https://github.com/soccin/BIC-variants_pipeline and the post processing code is at https://github.com/soccin/Variant-PostProcess.

### RNA-seq analysis

Raw sequencing reads were 3’ trimmed for quality <15 and adapters using version 0.4.5 of TrimGalore (https://www.bioinformatics.babraham.ac.uk/projects/trim_galore), and then aligned to mouse assembly mm9 with STAR v2.4 using default parameters. Post-alignment quality and transcript coverage were assessed using the Picard tool CollectRNASeqMetrics (http://broadinstitute.github.io/picard/). Raw read count tables were created using HTSeq v0.9.1. Normalization and expression dynamics were evaluated with DESeq2 using the default parameters including library size factor normalization. Heat maps were created using z-transformed normalized counts and plotted using pheatmap in R.

### Epigenomic Analysis

ATAC sequencing reads were 3’ trimmed and filtered for quality and adapter content using version 0.4.5 of TrimGalore, with a quality setting of 15, and running version 1.15 of cutadapt and version 0.11.5 of FastQC. Reads were aligned to mouse assembly mm9 with version 2.3.4.1 of bowtie2 (http://bowtie-bio.sourceforge.net/bowtie2/index.shtml) and were deduplicated using MarkDuplicates in version 2.16.0 of Picard Tools. To ascertain enriched regions, MACS2 (https://github.com/taoliu/MACS) was used with a p-value setting of 0.001. A global peak atlas was created by first removing blacklisted regions (http://mitra.stanford.edu/kundaje/akundaje/release/blacklists/mm9-mouse/mm9-blacklist.bed.gz), then defining a peak as +/- 250 bp around peak summits, and finally counting reads with version 1.6.1 of featureCounts (http://subread.sourceforge.net). Comparison of intra vs inter-group clustering in principle component analysis was used to determine normalization strategy, using either the median ratio method of DESeq2 or a sequencing depth-based factor normalized to ten million uniquely mapped fragments. The BEDTools suite (http://bedtools.readthedocs.io) was used to create normalized read density profiles based on the DESeq2 size factors. Differential enrichment was scored using DESeq2 for all pairwise group contrasts. All differential peaks were then merged for all contrasts in a given dataset, and k-means clustering was performed from k=4 to the point at which cluster groups became redundant. Peak-gene associations were created using linear genomic distance to transcription start site. GSEA for epigenomic data was performed with the pre-ranked option and default parameters, where each gene was assigned the single peak with the largest (in magnitude) log2 fold change associated with it. Motif signatures were obtained using Homer v4.5 (http://homer.ucsd.edu). Composite and tornado plots were created using deepTools v3.3.0 by running computeMatrix and plotHeatmap on normalized bigwigs with average signal sampled in 25 bp windows and flanking region defined by the surrounding 2 kb.

### Statistics

The statistical analysis was performed with GraphPad V10. Specific tests used as well as number of biological replicates is indicated in the legends corresponding to the respective figures.

### Study approval

All animal studies were reviewed and approved by the IACUC of Memorial Sloan Kettering Cancer Center, New York, New York, USA. For *in vivo* orthotopic mouse experiments we exclusively used female mice. For GEMM, male and female mice were used.

## Author contributions

Conceptualization including ideas and research aims: V.T., J.A.K. and J.A.F.

Experimental design and creation of models: V.T., J.J., G.P.K., J.A.K. and J.A.F.

Computational analysis: V.T. J.A.K., R.P.K. and N.D.S.

Data analysis V.T., J.D.F., B.X., R.G.

Data generation: V.T. J.G., T.Q., I.S.Y., J.A.K., L.B., E.J.S., A.L.H.

Manuscript preparation: V.T. and J.A.F.

Supervision: V.T., J.A.F.

## Acknowledgements

This study was supported in part by NIH grants CA255211, CA249663 and CA50706 to J.A.F. and from the Deutsche Forschungsgesellschaft (DFG) and The Mark Foundation for Cancer Research for V.T. We acknowledge the use of the following core facilities at Memorial Sloan Kettering Cancer Center, funded by the NCI Cancer Center Support Grant (CCSG, P30 CA08748): The Bioinformatics Core Facility, the Flow Cytometry Core Facility, the Molecular Cytology Core Facility, the Antitumor Assessment Core facility and the Integrated Genomics Operation (IGO) Core. IGO is additionally funded by Cycle for Survival and the Marie-Josée and Henry R. Kravis Center for Molecular Oncology. We are grateful to Pablo Sánchez Vela for help with figure design.

## Data availability

The data necessary to replicate the findings, except those in Figure 3A (Oncoprint from Human Specimens) will be made publicly available. Additionally, raw and normalized sequencing data from murine samples will be uploaded to GEO and made available upon publication.

