## Supplemental material for "Loss of tumor cell MHC Class II drives insensitivity of BRAF-mutant anaplastic thyroid cancers to MAPK inhibitors"

Conflict of interest statement: The authors declare no conflicts of interest with this work.

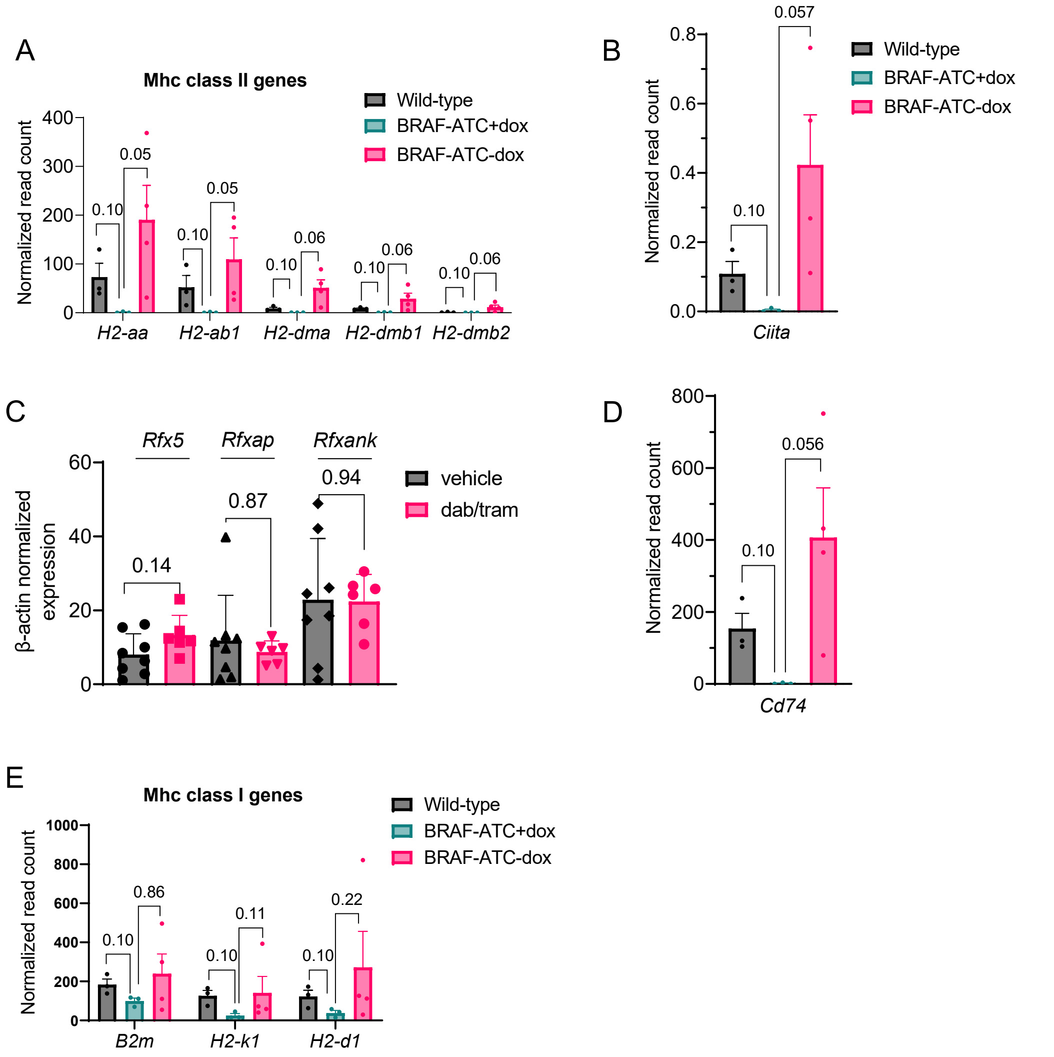


**Supplementary Figure 1**: A) Expression of MhcII complex genes is low or absent in WT thyrocytes, low in BRAF/p53-ATC cells and induced by dox withdrawal. B) *Ciita* expression in WT thyrocytes and in ATC cells prior to or after dox withdrawal. C) Quantitative RT-PCR of the MhcII-related transcription factors Rfx5, Rfxap and Rfxank is not impacted by dab/tram treatment in the orthotopic Braf/p53 model. D) *Cd74* expression in WT thyrocytes and in ATC cells prior to or after dox withdrawal. E) Expression of MhcI genes in WT thyrocytes or ATC cells prior to or after dox withdrawal. Multiple Mann-Whitney tests (A-E). Bars represent SEM. IFNγ: Interferon γ; GEMM: Genetic engineered mouse model; dab/tram: Dabrafenib and trametinib; Ciita: Class II major histocompatibility complex transactivator; WT: wild type; SEM: Standard error of the mean.

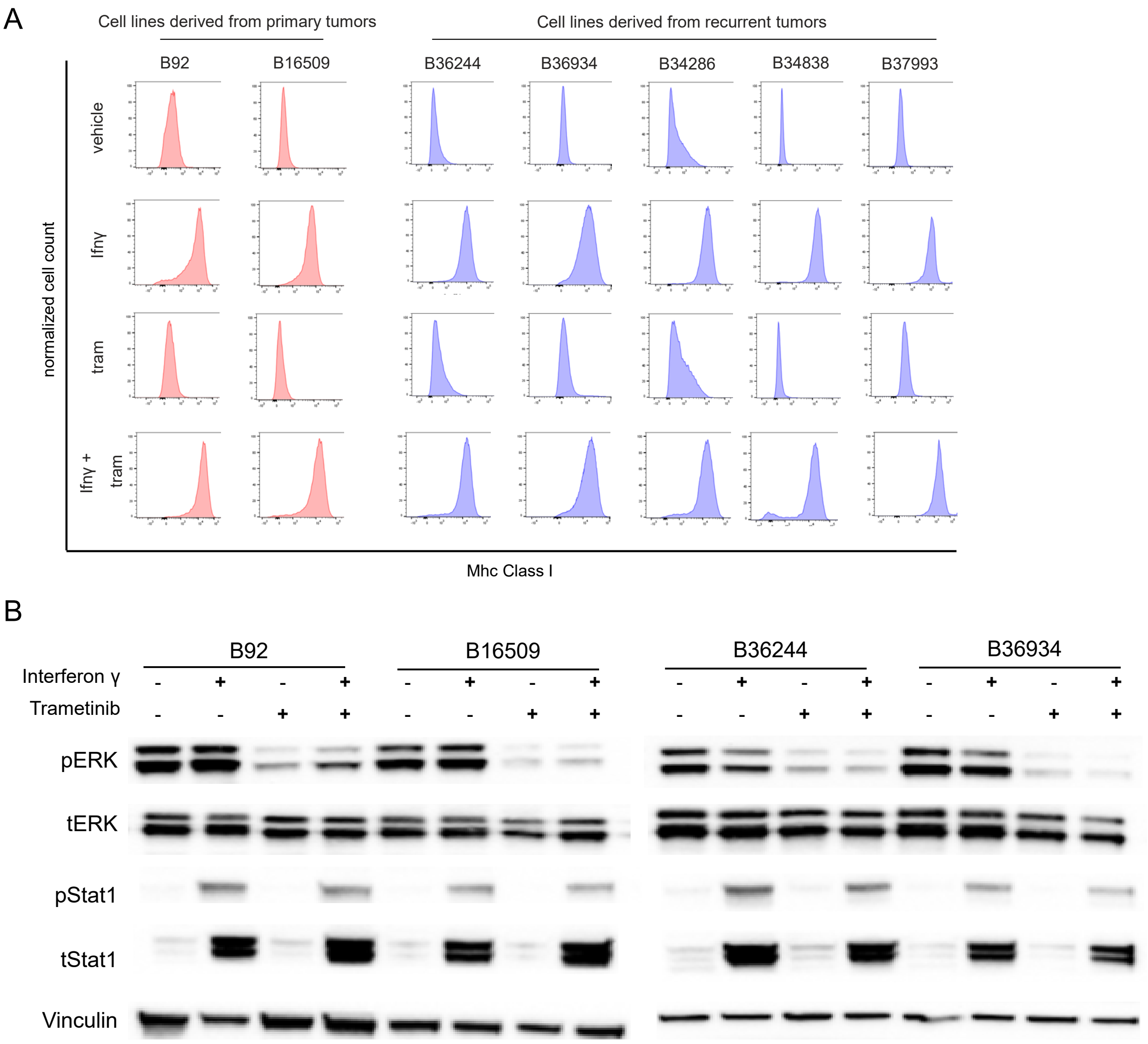


**Supplementary Figure 2**: A) Induction of MhcI by IFNγ (20ng/ml) as determined by FACS in two primary cell lines derived from primary ATCs on dox diet (B92 and B16509) and five lines derived from recurrent tumors developing after dox withdrawal (B36244, B36934, B34286, B34838 and B37993). Addition of trametinib (10nM) does not significantly augment the IFNγ (20ng/ml) effect. B) Western blot of B92, B16509, B36244, B36934 cell lysates treated with trametinib alone or in combination with IFNγ for 96 h for the indicated proteins. IFNγ: Interferon γ; dox: doxycycline.

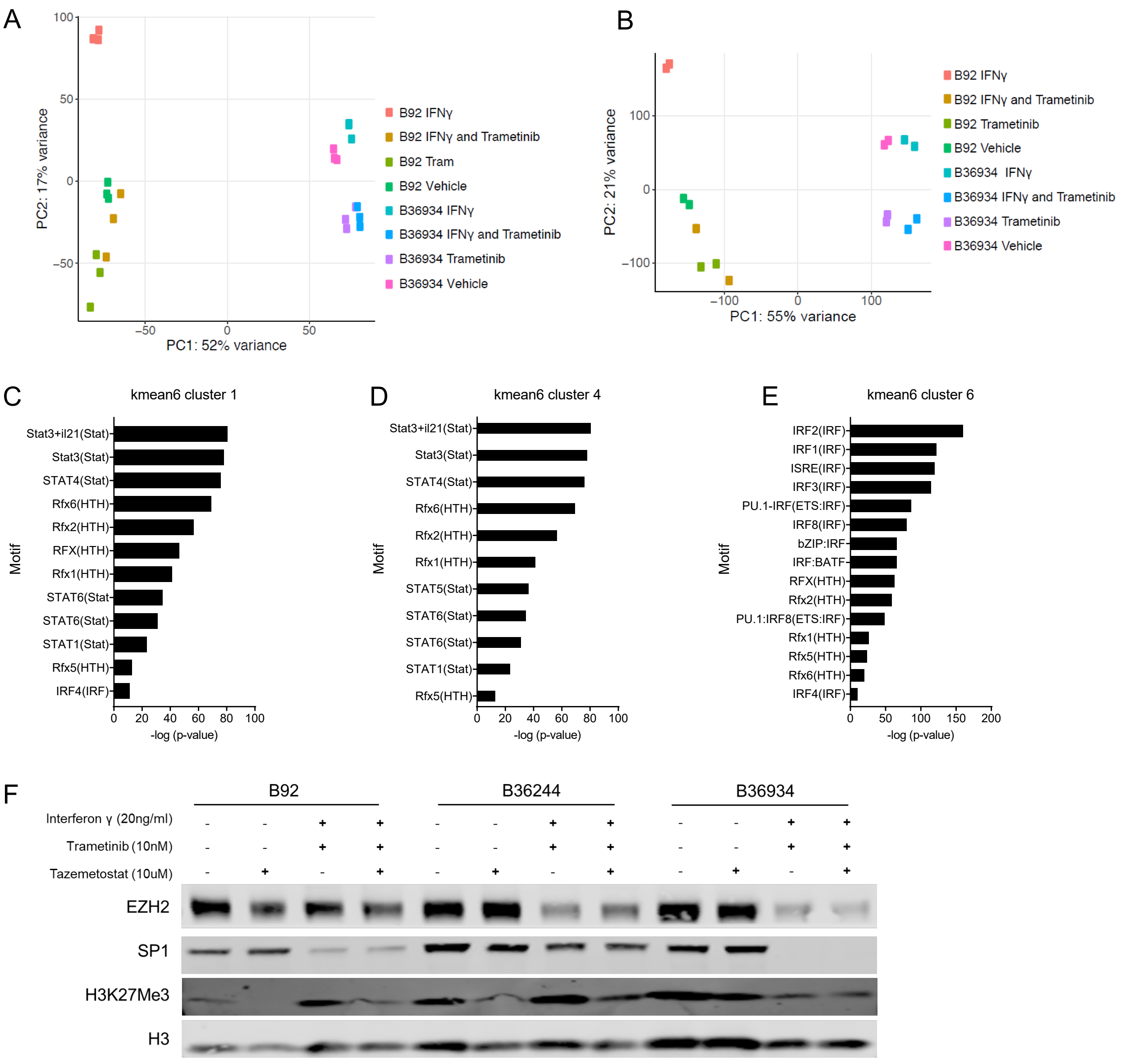


**Supplementary Figure 3:** A,B) Principal component analyses of RNA Seq and ATAC-Seq for the B92 primary and the B36934 recurrent ATC lines displaying triplicates and duplicates, respectively, for each treatment condition (DMSO, IFNγ (20ng/ml), trametinib (10nM) and the combination of IFNγ + trametinib (10nM) for 96 h. C–E) TF motifs of members of the Rfx, Stat and Irf families enriched in kmeans clusters 1, 4 and 6 identified using HOMER *de novo* motif discovery. F) Western blot for EZH2, Sp1, H3K27Me3 and H3 for the indicated cell lines and treatment conditions. IFNγ: Interferon γ; tram: trametinib; TF: transcription factor.

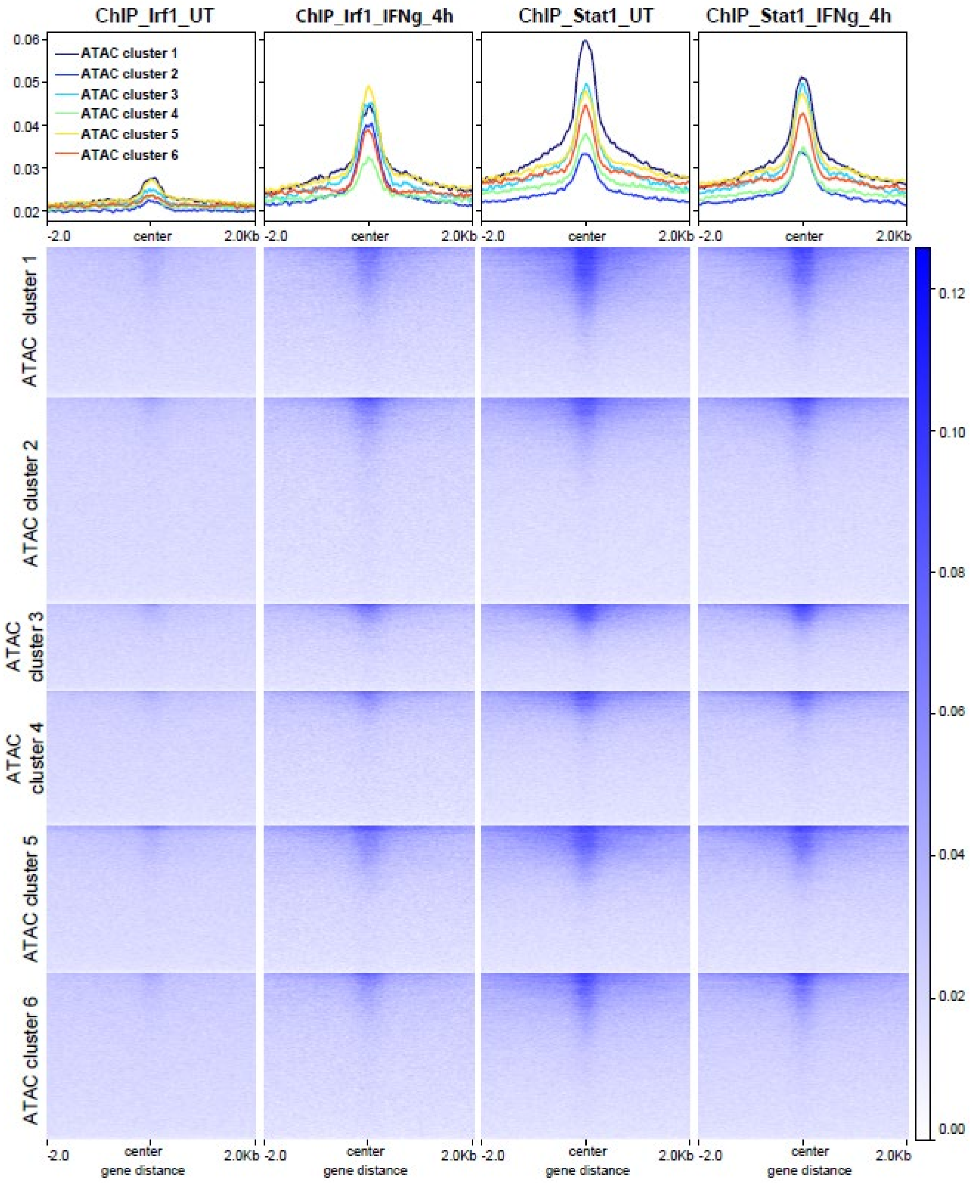


**Supplementary Figure 4**: Tornado plots of averaged Stat1 and Irf1 transcription factor binding sites in the respective k-means clusters.

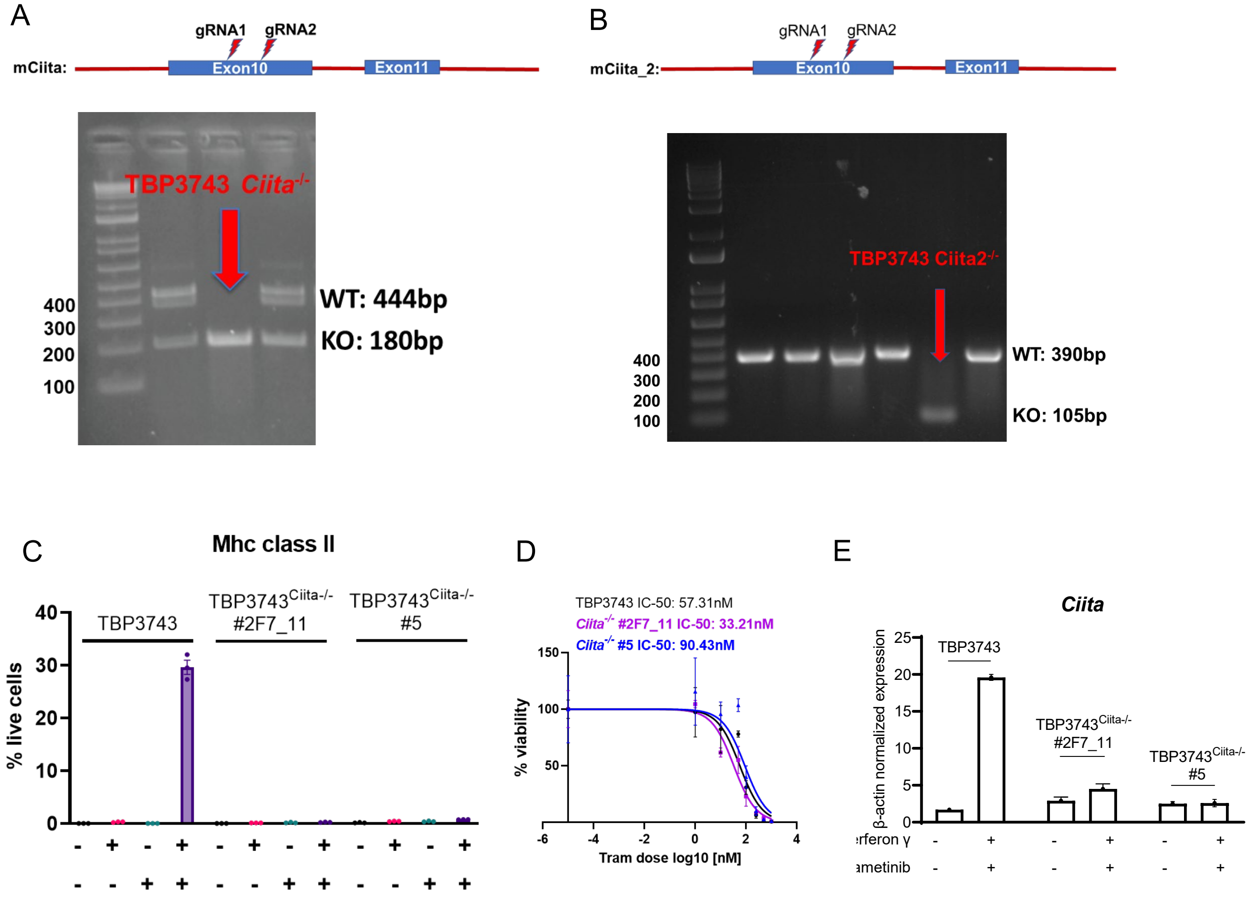


**Supplementary Figure 5**: A, B) *Top:* Schema of two different gRNA pairs targeting Exon 10 for CRISPR KO of *Ciita. Bottom:* Image of gel electrophoresis of PCR products from the indicated CRISPR KO clones. C) Loss of IFNγ and trametinib-induced MhcII in the *Ciita* CRISPR KO clones 2F7_11 and 5 derived from TBP3743 cells. MhcII was measured by FACS 96h after treatment with the indicated conditions. D) $IC_{50}$ for trametinib in parental TBP3743 cells and the *Ciita*$^{-/-}$ clones 2F7_11 and 5. E) Quantitative RT-PCR of Ciita mRNA in parental and *Ciita*$^{-/-}$ clones. gRNA: guide RNA; IFNγ: Interferon γ; Ciita: Class II major histocompatibility complex transactivator.

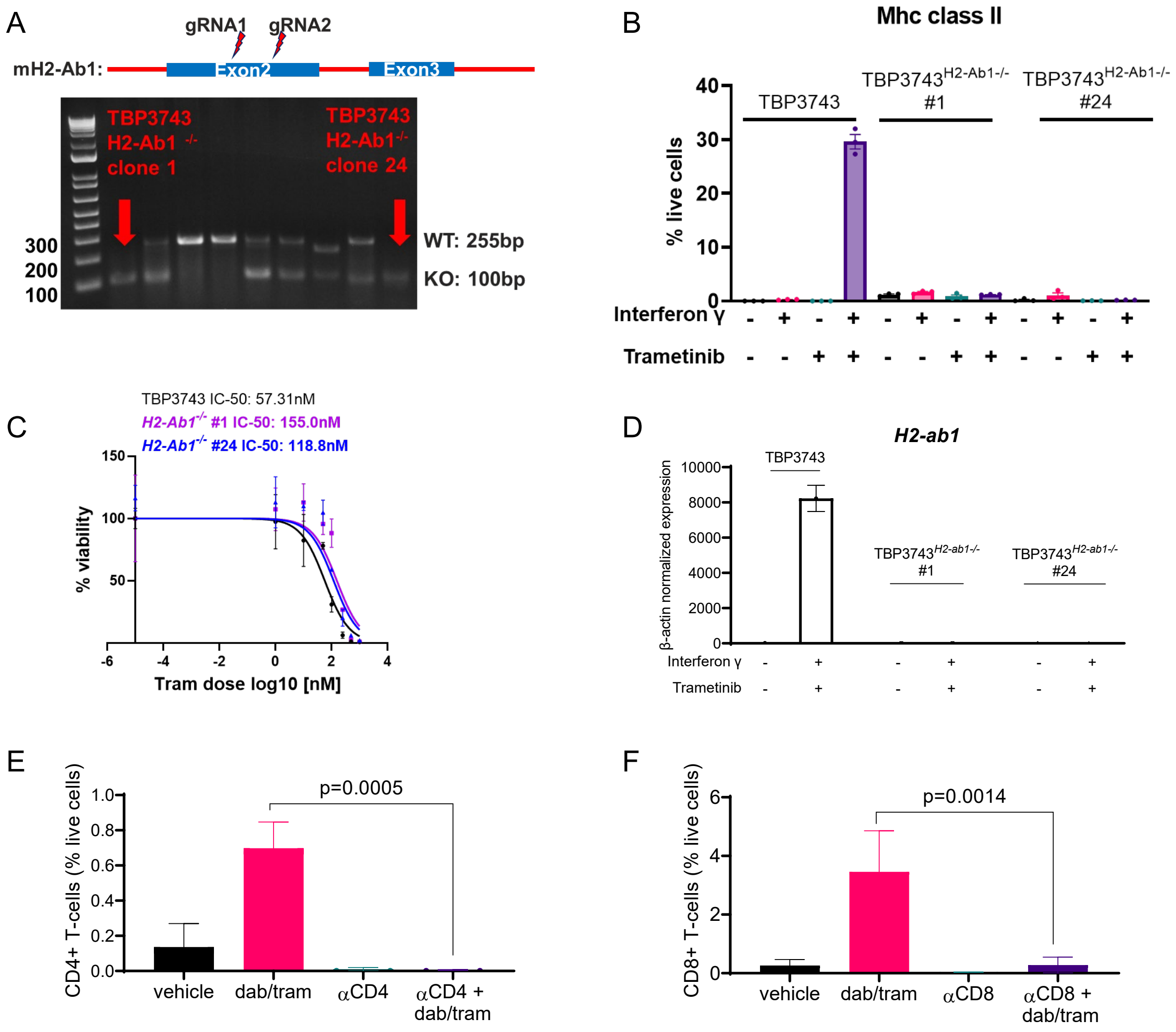


**Supplementary Figure 6:** A) *Top:* Schema for gRNA pair targeting exon2 of *H2-ab1* for generation of CRISPR KO clones. *Bottom:* Image of gel electrophoresis of PCR products of two homozygous *H2-ab1* KO clones. B) Loss of IFNγ + trametinib-induced MhcII in the *H2-ab1* CRISPR KO clones 1 and 24. Treatment conditions were as in Fig S4 C. C) $IC_{50}$ for trametinib in parental and *H2-ab1*$^{-/-}$ clones 1 and 24. D) Quantitative RT-PCR for H2-ab1 mRNA in parental and H2-ab1 CRISPR KO clones. E) and F) CD4+ (E) and CD8+ T-cells (G) percentage of live cells in thyroids harvested from mice treated with in the indication conditions. gRNA: guide RNA; IFNγ: Interferon γ; Ciita: Class II major histocompatibility complex transactivator.

Supplementary Tables

Table 1 Tumor mutational burden in human and murine papillary and anaplastic thyroid cancer
Table 2 Whole exome sequencing results: High confidence Maf
Table 3 Antibodies for IHC, IF, FACS and WB
Table 4 CrisprKO gRNA Seqences
Table 5 Primers

**Supplementary Table 1**

| | Human ATC | Murine ATC |
|---|---|---|
| Mutation per tumor | 23.7 [1] | 22 [3] |
| | 24 [2] | |

Supplementary Table 2

| Hugo_Symb | Entrez_ | Cente | NCBI_Build | Chromo | Start_ | End_Positio | Stran | Variant_Classificatio | Variant_ | Referenc | Tumor_Seq | Tumor_Seq | dbSNP_RS | Tumor_Sampl | Matched_Norm_ | Match_Norm_ |
|---|---|---|---|---|---|---|---|---|---|---|---|---|---|---|---|---|
| Grb14 | 0 | . | GRCm38 | 2 | 64930221 | 64930222 | + | Frame_Shift_Ins | INS | - | - | T | novel | s_B34286 | s_B34286_NT | - |
| [illegible] | 0 | [illegible] | GRCm38 | 2 | [illegible] | [illegible] | + | Missense_Mutation | SNP | T | T | C | novel | s_B34286 | s_B34286_NT | T |
| Olfr1156 | 0 | . | GRCm38 | 2 | 87949886 | 87949886 | + | Missense_Mutation | SNP | C | C | G | novel | s_B36244 | s_B36244_NT | C |
| Olfr1238 | 0 | . | GRCm38 | 2 | 89406938 | 89406938 | + | Missense_Mutation | SNP | A | A | G | novel | s_B36934 | s_B36934_NT | A |
| [illegible] | 0 | [illegible] | GRCm38 | 2 | 1.05E+08 | 1.05E+08 | + | Missense_Mutation | SNP | T | T | G | novel | s_B36244 | s_B36244_NT | T |
| [illegible] | 0 | . | GRCm38 | 2 | 1.5E+08 | 1.5E+08 | + | Missense_Mutation | SNP | [illegible] | [illegible] | [illegible] | novel | [illegible] | [illegible] | [illegible] |
| Slc7a11 | 0 | . | GRCm38 | 3 | 50442999 | 50442999 | + | Missense_Mutation | SNP | A | A | G | novel | s_B36934 | s_B36934_NT | A |
| [illegible] | 0 | [illegible] | GRCm38 | 3 | [illegible] | [illegible] | + | Missense_Mutation | SNP | T | T | G | novel | s_B36934 | s_B36934_NT | T |
| [illegible] | 0 | . | GRCm38 | [illegible] | [illegible] | [illegible] | + | Frame_Shift_Ins | INS | - | - | [illegible] | novel | [illegible] | [illegible] | - |
| Ankrd35 | 0 | . | GRCm38 | 3 | 96683851 | 96683852 | + | Frame_Shift_Ins | INS | - | - | A | novel | s_B36244 | s_B36244_NT | - |
| [illegible] | 0 | [illegible] | GRCm38 | 3 | 1.17E+08 | 1.17E+08 | + | Missense_Mutation | SNP | C | C | A | novel | s_B36934 | s_B36934_NT | C |
| [illegible] | 0 | . | GRCm38 | [illegible] | [illegible] | [illegible] | + | Missense_Mutation | SNP | [illegible] | [illegible] | [illegible] | novel | [illegible] | [illegible] | [illegible] |
| Cfi | 0 | . | GRCm38 | 3 | 1.3E+08 | 1.3E+08 | + | Missense_Mutation | SNP | C | C | T | rs30631084 | s_B36244 | s_B36244_NT | C |
| [illegible] | 0 | . | GRCm38 | 3 | 1.39E+08 | 1.39E+08 | + | Missense_Mutation | SNP | T | T | G | novel | s_B36934 | s_B36934_NT | T |
| [illegible] | 0 | . | GRCm38 | [illegible] | [illegible] | [illegible] | + | Missense_Mutation | SNP | [illegible] | [illegible] | [illegible] | novel | [illegible] | [illegible] | [illegible] |
| Cachd1 | 0 | . | GRCm38 | 4 | 1.01E+08 | 1.01E+08 | + | Missense_Mutation | SNP | A | A | G | novel | s_B34286 | s_B34286_NT | A |
| [illegible] | 0 | . | GRCm38 | 4 | 1.25E+08 | 1.25E+08 | + | Splice_Site | INS | - | - | C | novel | s_B34286 | s_B34286_NT | - |
| [illegible] | 0 | . | GRCm38 | 4 | [illegible] | [illegible] | + | Splice_Site | SNP | [illegible] | [illegible] | [illegible] | novel | [illegible] | [illegible] | [illegible] |
| [illegible] | 0 | . | GRCm38 | 5 | 6770990 | 6770990 | + | Missense_Mutation | SNP | C | C | A | novel | s_B36244 | s_B36244_NT | C |
| Kmt2c | 0 | . | GRCm38 | 5 | 25298974 | 25298974 | + | Frame_Shift_Del | DEL | C | C | - | novel | s_B34286 | s_B34286_NT | C |
| [illegible] | 0 | . | GRCm38 | 5 | [illegible] | [illegible] | + | Missense_Mutation | SNP | [illegible] | [illegible] | [illegible] | novel | [illegible] | [illegible] | [illegible] |
| Ociad2 | 0 | . | GRCm38 | 5 | 73323816 | 73323816 | + | Missense_Mutation | SNP | T | T | C | novel | s_B34286 | s_B34286_NT | T |
| Clock | 0 | . | GRCm38 | 5 | 76227043 | 76227045 | + | In_Frame_Del | DEL | GCT | GCT | - | rs25706717 | s_B36244 | s_B36244_NT | GCT |
| [illegible] | 0 | . | GRCm38 | 5 | 1.07E+08 | 1.07E+08 | + | Missense_Mutation | SNP | [illegible] | [illegible] | A | [illegible] | [illegible] | [illegible] | [illegible] |
| Brdt | 0 | . | GRCm38 | 5 | 1.07E+08 | 1.07E+08 | + | Missense_Mutation | SNP | A | A | G | novel | s_B34286 | s_B34286_NT | A |
| [illegible] | 0 | . | GRCm38 | 5 | 1.1E+08 | 1.1E+08 | + | Missense_Mutation | SNP | A | A | G | novel | s_B34286 | s_B34286_NT | A |
| [illegible] | 0 | [illegible] | GRCm38 | 5 | 1.38E+08 | 1.38E+08 | + | In_Frame_Del | DEL | GGA | GGA | - | [illegible] | s_B36244 | s_B36244_NT | GGA |
| Hsph1 | 0 | . | GRCm38 | 5 | 1.5E+08 | 1.5E+08 | + | Missense_Mutation | SNP | T | T | C | novel | s_B34286 | s_B34286_NT | T |
| Samd9l | 0 | . | GRCm38 | 6 | 3376305 | 3376305 | + | Missense_Mutation | SNP | T | T | G | novel | s_B36244 | s_B36244_NT | T |
| [illegible] | 0 | [illegible] | GRCm38 | 6 | 43081526 | 43081526 | + | Missense_Mutation | SNP | T | T | G | novel | s_B36244 | s_B36244_NT | T |

[illegible]

| DOMAINS | PUBMED | IMPACT | PIC | VARIANT_CLASS | TSL | HGVS_OF | GENE |
|---|---|---|---|---|---|---|---|
| PROSITE_profiles:PS50003.CDD:cd01259.PANTHER:PTHR11243.PANTHER:PTHR11243:SF22.Gene3D:2.30.29.30.Pfam:PF00169.SMART:SM00233.Superfamily:SSF50729 | | HIGH | 1 | insertion | 1 | | 1 |
| PANTHER:PTHR10037.PANTHER:PTHR10037:SF221.Gene3D:1.10.287.70.Pfam:PF00520.Superfamily:SSF81324 | | MODERATE | 1 | SNV | 1 | | 1 |
| PROSITE_profiles:PS50262.CDD:cd15410.PANTHER:PTHR26452:SF57.PANTHER:PTHR26452.PROSITE_patterns:PS00237.Gene3D:1.20.1070.10.Pfam:PF13853.Superfamily:SSF81321.Prints:PR00237.Transmembrane_helices:TMhelix | | MODERATE | 1 | SNV | 5 | | |
| PROSITE_profiles:PS50262.CDD:cd15939.PANTHER:PTHR26451:SF821.PANTHER:PTHR26451.Pfam:PF13853.Gene3D:1.20.1070.10.Superfamily:SSF81321.Prints:PR00237.Transmembrane_helices:TMhelix | | MODERATE | 1 | SNV | 5 | | |
| Pfam:PF05794.PANTHER:PTHR12832.PANTHER:PTHR12832:SF15 | | MODERATE | 1 | SNV | 5 | | |
| Gene3D:3.30.160.60.Pfam:PF00096.PROSITE_patterns:PS00028.PROSITE_profiles:PS50157.PANTHER:PTHR24377.PANTHER:PTHR24377.PANTHER:PTHR24377:SF818.SMART:SM00355.Superfamily:SSF57667 | | MODERATE | 1 | SNV | 1 | | |
| Gene3D:1.20.1740.10.Pfam:PF13520.PIRSF:PIRSF006060.PANTHER:PTHR11785.PANTHER:PTHR11785:SF323.TIGRFAM:TIGR00911.Transmembrane_helices:TMhelix | | MODERATE | 1 | SNV | 1 | | 1 |
| PANTHER:PTHR24403.PANTHER:PTHR24403:SF59 | | MODERATE | 1 | SNV | 1 | | 1 |
| PANTHER:PTHR24161.PANTHER:PTHR24161:SF21 | | HIGH | 1 | insertion | 1 | | 1 |
| PANTHER:PTHR24161.PANTHER:PTHR24161:SF21 | | HIGH | 1 | insertion | 1 | 1 | 1 |
| | | MODERATE | 1 | SNV | 1 | | 1 |
| PANTHER:PTHR44074.PANTHER:PTHR44074:SF2.Gene3D:3.30.70.20.Gene3D:3.50.50.60 | | MODERATE | 1 | SNV | 1 | | 1 |
| PANTHER:PTHR24253.PANTHER:PTHR24253:SF91 | | MODERATE | 1 | SNV | 1 | | 1 |
| PANTHER:PTHR21580:SF28.PANTHER:PTHR21580 | | MODERATE | 1 | SNV | 1 | | |
| PANTHER:PTHR12000.PANTHER:PTHR12000:SF1.Pfam:PF01650.PIRSF:PIRSF500148.PIRSF:PIRSF019663.Gene3D:3.40.50.1460.Prints:PR00776 | | MODERATE | 1 | SNV | 1 | | 1 |
| Gene3D:3.30.450.20.PANTHER:PTHR10166.PANTHER:PTHR10166:SF40.Superfamily:SSF103190.CDD:cd12913 | | MODERATE | 1 | SNV | 1 | | 1 |
| | | HIGH | 1 | insertion | 1 | | 1 |
| | | HIGH | 1 | SNV | 1 | | 1 |
| PANTHER:PTHR17614.PANTHER:PTHR17614:SF12.MobiDB_lite:mobidb-lite | | MODERATE | 1 | SNV | 5 | | |
| PANTHER:PTHR45888.PANTHER:PTHR45888:SF1.MobiDB_lite:mobidb-lite.MobiDB_lite:mobidb-lite.Low_complexity_(Seg):seg | | HIGH | 1 | deletion | 5 | | 1 |
| Low_complexity_(Seg):seg.PANTHER:PTHR22014.MobiDB_lite:mobidb-lite.MobiDB_lite:mobidb-lite.Coiled-coils_(Ncoils):Coil | | MODERATE | 1 | SNV | 5 | | 1 |
| PANTHER:PTHR13336.PANTHER:PTHR13336:SF2 | | MODERATE | 1 | SNV | 1 | | |
| Low_complexity_(Seg):seg.PANTHER:PTHR46055:SF2.PANTHER:PTHR46055.MobiDB_lite:mobidb-lite | | MODERATE | 1 | deletion | 2 | | 1 |
| PANTHER:PTHR22880.PANTHER:PTHR22880.PANTHER:PTHR22880:SF175.PANTHER:PTHR22880:SF175.MobiDB_lite:mobidb-lite.MobiDB_lite:mobidb-lite.Low_complexity_(Seg):seg | | MODERATE | 1 | SNV | 1 | | 1 |
| PANTHER:PTHR22880.PANTHER:PTHR22880.PANTHER:PTHR22880:SF175.PANTHER:PTHR22880:SF175.MobiDB_lite:mobidb-lite.MobiDB_lite:mobidb-lite.Low_complexity_(Seg):seg | | MODERATE | 1 | SNV | 1 | | 1 |
| PANTHER:PTHR18902.PANTHER:PTHR18902:SF26.MobiDB_lite:mobidb-lite.MobiDB_lite:mobidb-lite | | MODERATE | 1 | SNV | 5 | | 1 |
| PANTHER:PTHR14445.PANTHER:PTHR14445:SF37.MobiDB_lite:mobidb-lite.MobiDB_lite:mobidb-lite.Low_complexity_(Seg):seg | | MODERATE | 1 | deletion | 1 | 25 | |
| CDD:cd11739.PANTHER:PTHR45639.PANTHER:PTHR45639:SF2.Gene3D:3.30.420.40.Gene3D:3.30.30.30.Pfam:PF00012.Superfamily:SSF53067 | | MODERATE | 1 | SNV | 1 | | 1 |
| PANTHER:PTHR16155.PANTHER:PTHR16155:SF18 | | MODERATE | 1 | SNV | 1 | | 1 |
| | | MODERATE | 1 | SNV | 5 | | |
| PANTHER:PTHR10648:SF26.PANTHER:PTHR10648.Pfam:PF01179.Gene3D:2.70.98.20.Superfamily:SSF49998 | | HIGH | 1 | deletion | 1 | 4 | |
| | | HIGH | 1 | SNV | 1 | | 1 |
| Low_complexity_(Seg):seg.MobiDB_lite:mobidb-lite.MobiDB_lite:mobidb-lite.Coiled-coils_(Ncoils):Coil | | MODERATE | 1 | SNV | 1 | | |
| Low_complexity_(Seg):seg.Pfam:PF15761.MobiDB_lite:mobidb-lite.MobiDB_lite:mobidb-lite.Coiled-coils_(Ncoils):Coil | | MODERATE | 1 | SNV | 1 | | |
| PANTHER:PTHR36170.Coiled-coils_(Ncoils):Coil | | HIGH | 1 | SNV | 1 | | |
| CDD:cd00190.Gene3D:2.40.10.10.Gene3D:2.40.10.10.Pfam:PF00089.SMART:SM00020.Superfamily:SSF50494.PROSITE_profiles:PS50240.PANTHER:PTHR24271.PANTHER:PTHR24271:SF47 | | MODERATE | 1 | SNV | 1 | | |
| PANTHER:PTHR47014.MobiDB_lite:mobidb-lite.MobiDB_lite:mobidb-lite.Low_complexity_(Seg):seg | | MODERATE | 1 | deletion | 1 | | |
| | | HIGH | 1 | SNV | 1 | | 1 |
| Gene3D:2.30.30.40.Pfam:PF00018.PIRSF:PIRSF001741.PROSITE_profiles:PS50002.PANTHER:PTHR23122.PANTHER:PTHR23122:SF67.SMART:SM00326.Superfamily:SSF50044.CDD:cd12032 | | MODERATE | 1 | SNV | 1 | | 1 |
| PANTHER:PTHR21254.Superfamily:SSF49562 | | HIGH | 1 | SNV | 5 | | 1 |
| Gene3D:1.20.1070.10.Pfam:PF13853.Prints:PR00237.PROSITE_profiles:PS50262.PANTHER:PTHR26450.PANTHER:PTHR26450:SF72.Superfamily:SSF81321.Transmembrane_helices:TMhelix.CDD:cd15222 | | MODERATE | | SNV | 5 | | |
| CDD:cd08403.PANTHER:PTHR10024:SF180.PANTHER:PTHR10024.Gene3D:2.60.40.150.SMART:SM00239.Superfamily:SSF49562 | | MODERATE | 1 | SNV | 1 | | |
| PANTHER:PTHR11905.PANTHER:PTHR11905:SF118.MobiDB_lite:mobidb-lite.MobiDB_lite:mobidb-lite | 2E+07 | MODERATE | 1 | SNV | 1 | | 1 |
| PANTHER:PTHR11905.PANTHER:PTHR11905:SF118.MobiDB_lite:mobidb-lite.MobiDB_lite:mobidb-lite | 2E+07 | MODERATE | 1 | SNV | 1 | | 1 |
| Gene3D:1.20.1070.10.Pfam:PF13853.Prints:PR00245.PROSITE_profiles:PS50262.PANTHER:PTHR24248.PANTHER:PTHR24248:SF71.Superfamily:SSF81321.Transmembrane_helices:TMhelix.CDD:cd15406 | | MODERATE | 1 | SNV | | | |
| Gene3D:3.40.70.480.Pfam:PF02147.PROSITE_profiles:PS51447.SMART:SM00896.Superfamily:SSF49991 | | MODERATE | 1 | SNV | 1 | | 1 |
| PANTHER:PTHR11584:SF332.PANTHER:PTHR11584 | | MODERATE | 1 | SNV | 1 | | 1 |
| Gene3D:2.10.25.10.Pfam:PF00053.PROSITE_profiles:PS50027.PANTHER:PTHR10574.PANTHER:PTHR10574:SF291.SMART:SM00180.SMART:SM00181.Superfamily:SSF57196.CDD:cd00055 | | MODERATE | 1 | SNV | 5 | | 1 |
| PANTHER:PTHR15742.PANTHER:PTHR15742:SF2 | | MODERATE | | SNV | 5 | | |
| Gene3D:3.30.70.330.Pfam:PF00076.PROSITE_profiles:PS50102.PANTHER:PTHR11176.PANTHER:PTHR11176:SF13.SMART:SM00360.Superfamily:SSF54928.CDD:cd12327 | | MODERATE | 1 | SNV | 1 | | 1 |
| Gene3D:3.80.10.10.Pfam:PF13855.PANTHER:PTHR45712.PANTHER:PTHR45712:SF13.SMART:SM00364.SMART:SM00369.Superfamily:SSF52058.Low_complexity_(Seg):seg | | MODERATE | 1 | SNV | 1 | | 1 |
| | | HIGH | 1 | SNV | 1 | | 1 |
| PANTHER:PTHR12874:SF17.PANTHER:PTHR12874 | | MODERATE | 1 | SNV | 1 | | |
| PANTHER:PTHR45620.PANTHER:PTHR45620:SF23.MobiDB_lite:mobidb-lite | | MODERATE | 1 | SNV | 1 | | 1 |
| PANTHER:PTHR34439.Coiled-coils_(Ncoils):Coil | | MODERATE | 1 | SNV | 1 | | |
| Gene3D:2.60.40.820.Pfam:PF00907.PROSITE_profiles:PS50252.PANTHER:PTHR11267.PANTHER:PTHR11267:SF125.SMART:SM00425.Superfamily:SSF49417.CDD:cd00182 | | MODERATE | 1 | SNV | 1 | | 1 |
| PANTHER:PTHR24361.PANTHER:PTHR24361:SF651.PIRSF:PIRSF038175 | | MODERATE | 1 | SNV | 1 | | 1 |
| PANTHER:PTHR22656 | | MODERATE | | SNV | 5 | | |
| PROSITE_profiles:PS50157.PANTHER:PTHR24394:SF51.PANTHER:PTHR24394:SF51.PANTHER:PTHR24394.PANTHER:PTHR24394.Gene3D:3.30.160.60.Superfamily:SSF57667 | | MODERATE | 1 | SNV | 5 | | |
| PANTHER:PTHR15036.PANTHER:PTHR15036:SF15 | | MODERATE | 1 | SNV | 5 | | |
| PROSITE_profiles:PS50262.CDD:cd15913.PANTHER:PTHR24242:SF239.PANTHER:PTHR24242.Gene3D:1.20.1070.10.Pfam:PF13853.Superfamily:SSF81321.Prints:PR00245 | | MODERATE | 1 | SNV | 5 | | |
| Gene3D:1.10.630.10.Pfam:PF00067.Prints:PR00385.PANTHER:PTHR24300.PANTHER:PTHR24300:SF119.Superfamily:SSF48264.Transmembrane_helices:TMhelix.Low_complexity_(Seg):seg | | MODERATE | 1 | SNV | 1 | | |
| PANTHER:PTHR21584:SF10.PANTHER:PTHR21584.MobiDB_lite:mobidb-lite | | MODERATE | | SNV | | | |
| Pfam:PF00083.PROSITE_profiles:PS50850.PANTHER:PTHR23500.PANTHER:PTHR23500:SF198.Superfamily:SSF103473.TIGRFAM:TIGR00879.CDD:cd17360 | | MODERATE | 1 | SNV | 1 | | 1 |
| Gene3D:1.20.120.450.Pfam:PF00520.PANTHER:PTHR10037.PANTHER:PTHR10037:SF227.Superfamily:SSF81324.Transmembrane_helices:TMhelix | | MODERATE | 1 | SNV | 5 | | 1 |
| Gene3D:3.40.50.300.Pfam:PF00009.PROSITE_profiles:PS51722.PANTHER:PTHR23115.PANTHER:PTHR23115:SF125.Superfamily:SSF52540.CDD:cd01883 | | MODERATE | 1 | SNV | 1 | | |
| PANTHER:PTHR10507:SF0.PANTHER:PTHR10507.Pfam:PF02724 | | MODERATE | 1 | SNV | 1 | | 1 |
| PANTHER:PTHR10507:SF0.PANTHER:PTHR10507.Pfam:PF02724 | | HIGH | 1 | SNV | 1 | | 1 |
| | | HIGH | 1 | SNV | 1 | | |
| MobiDB_lite:mobidb-lite.Low_complexity_(Seg):seg | | MODERATE | 1 | SNV | 5 | | 1 |
| Gene3D:2.60.40.10.PROSITE_profiles:PS50853.PANTHER:PTHR12231.PANTHER:PTHR12231:SF242.Superfamily:SSF49265 | | MODERATE | 1 | SNV | 1 | | 1 |
| Gene3D:2.60.40.10.PANTHER:PTHR11412.PANTHER:PTHR11412:SF86 | | HIGH | 1 | SNV | 1 | | 1 |
| Superfamily:SSF81321.PANTHER:PTHR26453:SF64.PANTHER:PTHR26453 | | MODERATE | 1 | SNV | 5 | | |
| PANTHER:PTHR45874.PANTHER:PTHR45874:SF1.PIRSF:PIRSF036604 | | HIGH | 1 | insertion | 1 | | 1 |
| PANTHER:PTHR28602.MobiDB_lite:mobidb-lite.Low_complexity_(Seg):seg | | MODERATE | 1 | deletion | 1 | | 1 |
| PROSITE_profiles:PS50020.CDD:cd00201.PANTHER:PTHR15377.PROSITE_patterns:PS01159.Pfam:PF00397.Gene3D:2.20.70.10.SMART:SM00456.Superfamily:SSF51045 | | MODERATE | 1 | SNV | | | 1 |
| Gene3D:1.10.510.10.Pfam:PF16474.PROSITE_profiles:PS51377.PANTHER:PTHR21345.PANTHER:PTHR21345:SF8.SMART:SM00750.Low_complexity_(Seg):seg | | MODERATE | 1 | deletion | 1 | | 1 |
| Pfam:PF04758.PANTHER:PTHR12650.PANTHER:PTHR12650:SF15.MobiDB_lite:mobidb-lite.MobiDB_lite:mobidb-lite.Low_complexity_(Seg):seg | | HIGH | 1 | insertion | 1 | | |
| Pfam:PF04758.PANTHER:PTHR12650.PANTHER:PTHR12650:SF15.MobiDB_lite:mobidb-lite.MobiDB_lite:mobidb-lite.Low_complexity_(Seg):seg | | HIGH | 1 | insertion | 1 | | |
| Pfam:PF04758.PANTHER:PTHR12650.PANTHER:PTHR12650:SF15.MobiDB_lite:mobidb-lite.MobiDB_lite:mobidb-lite.Low_complexity_(Seg):seg | | HIGH | 1 | insertion | 1 | | |
| Pfam:PF04758.PANTHER:PTHR12650.PANTHER:PTHR12650:SF15.MobiDB_lite:mobidb-lite.MobiDB_lite:mobidb-lite.Low_complexity_(Seg):seg | | MODERATE | 1 | deletion | 1 | | |
| Pfam:PF04758.PANTHER:PTHR12650.PANTHER:PTHR12650:SF15.MobiDB_lite:mobidb-lite.MobiDB_lite:mobidb-lite.Low_complexity_(Seg):seg | | HIGH | 1 | deletion | 1 | | |
| Pfam:PF04758.PANTHER:PTHR12650.PANTHER:PTHR12650:SF15 | | HIGH | 1 | insertion | 1 | 1 | |
| Pfam:PF04758.PANTHER:PTHR12650.PANTHER:PTHR12650:SF15 | | HIGH | 1 | insertion | 1 | | |
| Pfam:PF04758.PANTHER:PTHR12650.PANTHER:PTHR12650:SF15 | | HIGH | 1 | insertion | 1 | | |
| PANTHER:PTHR13190.PANTHER:PTHR13190:SF21 | | MODERATE | 1 | SNV | 1 | | |
| CDD:cd15407.PANTHER:PTHR26452.PANTHER:PTHR26452:SF482.Pfam:PF13853.Gene3D:1.20.1070.10.Superfamily:SSF81321.Prints:PR00237 | | MODERATE | 1 | SNV | 2 | | |
| PROSITE_profiles:PS50262.CDD:cd15225.PANTHER:PTHR26452.PANTHER:PTHR26452:SF165.Gene3D:1.20.1070.10.Pfam:PF13853.Superfamily:SSF81321.Prints:PR00245.Prints:PR00237.Transmembrane_helices:TMhelix | | MODERATE | 1 | SNV | 5 | | |
| | | HIGH | 1 | insertion | 1 | | 1 |
| PANTHER:PTHR41095 | | MODERATE | 1 | SNV | 5 | | |
| PANTHER:PTHR11219.PANTHER:PTHR11219:SF7.Gene3D:2.120.10.30.Superfamily:SSF101898 | | MODERATE | 1 | SNV | 1 | | |
| Pfam:PF02990.PANTHER:PTHR10766.PANTHER:PTHR10766:SF43.CDD:cd17489 | | MODERATE | 1 | SNV | 2 | | |
| PIRSF:PIRSF000254.PANTHER:PTHR24543.PANTHER:PTHR24543:SF249 | | MODERATE | 1 | SNV | 1 | | 1 |
| PANTHER:PTHR12317.PANTHER:PTHR12317:SF19.Pfam:PF03982.Superfamily:SSF69593 | | MODERATE | 1 | SNV | 5 | | 1 |
| Pfam:PF01391.PANTHER:PTHR24023.PANTHER:PTHR24023:SF877 | | MODERATE | 1 | SNV | 1 | | 1 |
| PDB-ENSP_mappings:5a72.M.PDB-ENSP_mappings:5a73.M.PANTHER:PTHR44507.PANTHER:PTHR44507:SF1.Pfam:PF00361.TIGRFAM:TIGR01972.Prints:PR01437.Transmembrane_helices:TMhelix | | MODERATE | 1 | SNV | | | |

**Supplementary Table 3**

| Antibody | Clone | Source | Target Species |
|---|---|---|---|
| HLA-DR | TAL 1B5 | Santa Cruz | human |
| Pan Cytokeratin | na | Thermo Fisher | human |
| CD8 | 4B11 | Thermo Fisher | human |
| CD3 | SP7 | Abcam | human |
| CD163 | 10D6 | Abcam | human |
| CD68 | KP1 | Abcam | human |
| CD15 | MMA | Abcam | human |
| BUV395 Rat Anti-Mouse CD45 | 30-F11 | BD Biosciences | mouse |
| BV510 Rat Anti-CD11b | M1/70 | BD Biosciences | mouse |
| BV750 Rat Anti-Mouse Siglec-F | E50-2440 | BD Biosciences | mouse |
| FITC Rat Anti-Mouse CD3 | 17A2 | BD Biosciences | mouse |
| BUV805 Rat Anti-Mouse CD8a | 53-6.7 | BD Biosciences | mouse |
| BUV661 Mouse Anti-Mouse NK-1.1 | PK136 | BD Biosciences | mouse |
| PerCP anti-mouse Ly-6G Antibody | 1A8 | Biolegend | mouse |
| Brilliant Violet 711™ anti-mouse CD11c Antibody | N418 | Biolegend | mouse |
| Spark Blue™ 550 anti-mouse I-A/I-E Antibody | M5/114.15.2 | Biolegend | mouse |
| Brilliant Violet 650™ anti-mouse F4/80 Antibody | BM8 | Biolegend | mouse |
| Brilliant Violet 570™ anti-mouse CD4 Antibody | RM4-5 | Biolegend | mouse |
| Spark NIR™ 685 anti-mouse/human CD45R/B220 Antibody | RA3-6B2 | Biolegend | mouse |
| APC-Cy™7 Rat Anti-Mouse Ly-6C | AL-21 | BD Biosciences | mouse |
| Arginase 1 Monoclonal Antibody PerCP-eFluor™ 710[1] | A1exF5 | Thermo Fisher | mouse |
| FOXP3 Monoclonal Antibody eFluor™ 450[1] | FJK-16s | Thermo Fisher | mouse |
| CD274 (PD-L1, B7-H1) Super Bright™ 436 | MIH5 | Thermo Fisher | mouse |
| BV605 Rat Anti-Mouse I-A/I-E | M5/114.15.2 | BD Biosciences | mouse |
| LIVE/DEAD™ Fixable Blue Dead Cell Stain Kit, for UV excitation | na | Thermo Fisher | mouse |
| Alexa Fluor® 647 anti-mouse CD206[1] | MMR | Biolegend | mouse |
| PE Mouse Anti-Mouse H-2Kb | AF6-88.5 | BD Biosciences | mouse |
| Phospho-p44/42 MAPK (Erk1/2) (Thr202/Tyr204) (D13.14.4E) XP® Rabbit mAb | | Cell Signaling | |
| p44/42 MAPK (Erk1/2) (L34F12) Mouse mAb | | Cell Signaling | |
| Phospho-Stat1 (Ser727) (D3B7) Rabbit mAb | | Cell Signaling | |
| Stat1 (D1K9Y) Rabbit mAb | | Cell Signaling | |
| Vinculin (E1E9V) XP® Rabbit mAb | | Cell Signaling | |
| SP1 (D4C3) Rabbit mAb | | Cell Signaling | |
| Ezh2 (D2C9) XP® Rabbit mAb | | Cell Signaling | |
| Histone H3 (D1H2) XP® Rabbit mAb | | Cell Signaling | |
| Tri-Methyl-Histone H3 (Lys27) (C36B11) Rabbit mAb | | Cell Signaling | |

[1] Intracellular antibodies

Supplementary Table 4

| Vector name | gRNA Sequence | Source | Identifier |
|---|---|---|---|
| mCiita_1 | AGCAGGCCAAGACTTACATG, TAGTCGAGCTGGCCAAGCTG | Vectorbuilder | VB210324-1175xee |
| mCiita_2 | CCCGGAGCCTTAGTCGAGCT, GAGACCCTATGACAACTGG | Vectorbuilder | VB230124-1305eyk |
| mH2-ab1 | ACGGGACGCAGCGCATACGA, GGAGATCCTGGAGCGAACGC | Vectorbuilder | VB230124-1307xfy |

Supplementary Table 5

| Primer | Sequence |
|---|---|
| mCiita_Ex3F | ATCTTCCAGCGGAAGCTACTGC |
| mCiita_Ex3R | CCGGGTTTCTTGCAAGGTGC |
| mCiita_Ex10F | GGACTCTATGTCAGCCTGCTAGG |
| mCiita_Ex10R | TGGGCTCGAGGCTGGAAAAC |
| mH2-ab1_Ex2F | CCGCAGGGCATTTCGTGTAC |
| mH2-ab1_Ex2R | TCTCCGGCCCCTCGTAGTTGT |
| mCD74_FW | AAGCAGTGGCTCTTGTTTGAG |
| mCD74_RV | CTTCCATGTCCAGTGGCTCT |
| mCiita_FW | AATCTACCACGGTGAGATGCCC |
| mCiita_RV | TCGGGGAGACTGGGGATACTGA |
| mH2-aa_FW | GAGCAGCTTCAGAGACCTCC |
| mH2-aa_RV | CTACGTGGTCGGCCTCAAT |
| mH2-ab1_FW | CACAGGAGTCAGAAAGGACCTC |
| mH2-ab1_RV | TGGCAGTCAGGAATTCGGAG |
| mH2-dma_FW | GAGATTGACCGCTACACGGCAA |
| mH2-dma_RV | GAAGACAATGCCCATGATGGTGC |
| mH2-dmb1_FW | AGAGCCTTCTCCAGCGTTTGCA |
| mH2-dmb1_RV | TGTGGTTTGGGCTACTCGGACA |
| mH2-dmb2_FW | ACCTTTCTGGGATGTGCTGACC |
| mH2-dmb2_RV | GTGATGGTCACATCCGCTGGAT |
| mRfxap_FW | ACGTCAAACTGGAGGAAAGCAC |
| mRfxap_RV | GAGTAGGTCTTGCAGGGCG |
| mRfx5_FW | GGAAGACCTTGGTATCCATGCC |
| mRfx5_RV | GGCTGCTTCTACCAGTTCATCC |
| Rfxank_FW | GCACATGCCTGTCTGGAAAC |
| Rfxank_RV | AGCAGGAAGCGAACTGTCTC |
| Irf1_FW | CAAAGCCACCATGCCAATCACTCG |
| Irf1_RV | GGCCCAGCTCCGGAACAGACAG |
| mH2-k1_FW | GGAGCAGGAGGGGCCCGAGTATTG |
| mH2-k1_RV | CGCCGTCCACGTTTTCAGGTCTTC |
| mB2m_FW | TCACTGACCGGCCTGTATGCTATC |
| mB2m_RV | AATGTGAGGCGGGTGGAACTGT |
